# Temporally organized representations of reward and risk in the human brain

**DOI:** 10.1101/2023.05.09.539916

**Authors:** Vincent Man, Jeffrey Cockburn, Oliver Flouty, Phillip E. Gander, Masahiro Sawada, Christopher K. Kovach, Hiroto Kawasaki, Hiroyuki Oya, Matthew A. Howard, John P. O’Doherty

## Abstract

The value and uncertainty associated with choice alternatives constitute critical features along which decisions are made. While the neural substrates supporting reward and risk processing have been investigated, the temporal organization by which these computations are encoded remains elusive. Here we leverage the high spatiotemporal precision of intracranial electroencephalography (iEEG) to uncover how representations of decision-related computations unfold in time. We present evidence of locally distributed representations of reward and risk variables that are temporally organized across multiple regions of interest. Reward outcome representations across wide-spread regions follow a temporally cascading order along the anteroposterior axis of the brain. In contrast, expected value can be decoded from multiple regions at the same time, and error signals in both reward and risk domains reflect a mixture of sequential and parallel encoding. We highlight the role of the anterior insula in generalizing between reward prediction error (RePE) and risk prediction error (RiPE), within which the encoding of RePE in the distributed iEEG signal predicts RiPE. Together our results emphasize the utility of uncovering temporal dynamics in the human brain for understanding how computational processes critical for value-based decisions under uncertainty unfold.

## 1 Introduction

Adaptive behaviour is predicated on an ability to evaluate predictive features of the environment. A substantial collection of evidence has demonstrated that humans and animals alike direct their actions according to reward expectations [1]. However, developing robust estimates of future reward is complicated by the fact that real-world settings are often inherently unpredictable. To act well in face of this unpredictability, the brain tracks and employs higher-order variables such as expected uncertainty (i.e. risk) [2, 3, 4, 5, 6]. Nevertheless, the temporal dynamics by which these variables are computed in the brain remain poorly described, largely due to the field’s reliance non-invasive neuroimaging techniques that are spatially or temporally coarse. To address this limitation, we leverage intracranial EEG (iEEG) to probe the neural evolution of reward and risk computations at high spatial and temporal resolution.

Reward prediction error (RePE) signals that quantify the discrepancy between observed and expected reward [7, 8] not only offer a means of iteratively improving reward estimates [9], but importantly also provide a quantity to describe outcome variability (i.e. risk) [10]. Analogous to RePE, it has been suggested that risk prediction error (RiPE) signals are computed by the brain when predictive associations between choice and outcome change [11, 5]. To explicitly investigate relationships between RePE and RiPE, we employ a normative model following from [12, 4] that describes RiPE as a second-order uncertainty term which is derived as a function of RePE, therefore making an explicit temporal prediction that neural signals reflecting reward error should precede and predict those supporting risk error computations.

However, little is known about the temporal characteristics of reward and risk representations at the wide spatial scale for which correlates of these decision variables have been reported in the human brain [13, 14]. Studies using scalp electroencephalography (EEG) from us [15] and others [16, 17] have begun to probe the neural correlates of reward and risk at finer timescales, with evidence suggesting a mixture of both parallel and sequential computations across the brain [15]. In support of this, an emerging line of work has begun to describe possible modes of temporal configuration by which cognitive processes are carried out in the brain. For example, evidence suggests that both parallel and sequential temporal arrangements support the encoding of visual features and their subsequent maintenance, respectively [18], and similar interactions between parallel and sequential processing modes have been described in the context of multi-tasking [19, 20]. Evidence of sequential processing in the reward domain has also highlighted a posterior to anterior flow of information [15, 21]. However, studies using scalp EEG provide limited spatial resolution and are unable to probe deeper subcortical regions [22], thereby painting an incomplete spatio-temporal mapping between theoretically grounded variables and neural function.

We aimed to expand our understanding of the temporal dynamics of reward processing in the brain by directly testing for interactions in the neural representations of pertinent variables. We investigate two theoretically motivated hypotheses; that RePE signals should precede risk prediction error, and that RePE representations should directly relate to risk error computations whereby the RePE serves as input to the computational function that defines risk error (see 4.8 for details). Importantly, evidence of an explicit correspondence between neural representations of different variables in a given brain region would lend support for that region’s role in supporting the computational transformations mid-stream. This hypothesis stands in contrast with the alternative possibility of distinct representational codes for different computational variables in the same neural population (e.g., [23]).

We also held strong prior hypotheses about candidate regions that would represent reward and risk variables. Within the prefrontal cortex, anterior and ventromedial PFC (vmPFC) have differential contributions to risk [24, 25] and reward [26, 27], respectively [28]. We further hypothesized that the amygdala would be important given its embedding in a reward circuit along with vmPFC [29] and OFC [30], and by our own previous work in human electrophysiology highlighting its role in reward representations [31, 32]. We further predicted that OFC and anterior insula in particular would be candidate substrates for holding representations across reward and risk domains. However, the temporal ordering of computational representations in OFC remains to be clarified given evidence of both simultaneous [33] and sequential [34] encoding of expected value and risk. Similarly, despite evidence of RePE [35, 36] and RiPE [17, 4, 37] signals in anterior insula, to date no studies have directly compared the relative timings nor tested for interactions between these variables in anterior insula.

In this pursuit, we recorded the human iEEG in a sample of patients undergoing chronic evaluation for epilepsy and implantation sites were determined on the basis of clinical considerations. Patients performed a task designed to decouple the rapid unfolding of both reward and risk variables first described by [12, 4], allowing us to investigate the neural mechanisms by which reward-related representations contribute to building risk computations. iEEG is well situated to overcome many of the spatiotemporal limtiations of noninvasive neuroimaging [38] and has been used to elucidate functional contributions of neural activity both within specific regions (e.g. [33, 39]) and across wide-spread areas of the brain [40, 41]. We leveraged this latter advantage of iEEG by simultaneously recording from frontal, subcortical and parietal areas to investigate the temporal organization of reward and risk encoding across widespread brain networks. Among the locations from which we were able to record the intracranial potential, we focused *a priori* on a smaller set of hypothesized ROIs while also exploring a wider span of regional targets. Our objectives were to address outstanding questions about the temporal signature of reward and risk encoding at scales unattainable with non-invasive human neuroimaging, and to test directly for neural interactions between computations.

We report wide-spread outcome processing that unfolds along the anteroposterior axis of the brain. In contrast, expected value can be decoded from multiple regions simultaneously, whereas error signals emerge through a mixture of sequential and parallel processing. We highlight the computational contribution of anterior insula in generalizing the RePE and risk prediction error (RiPE), where RePE signals reflected in the iEEG signal predicts subsequent RiPE signals. Together our results emphasize the utility of uncovering temporal dynamics in the human brain for understanding how reward and risk computations unfold.

## 2 Results

Participants (n=10) completed total of 16 recordings sessions in which we administered a simple gambling paradigm adapted from [12] (Figure 1A). On a trial, participants guessed whether the second of two sequentially-drawn playing cards would have a higher or lower numerical value than the first. After the second card was presented, participants had all the necessary information to determine the accuracy of their initial guess. Correct guesses earned game points and incorrect guesses resulted in point deductions. After each trial, participants reported whether they had won. The deck was shuffled and cards were replaced between trials.

**Figure 1:**
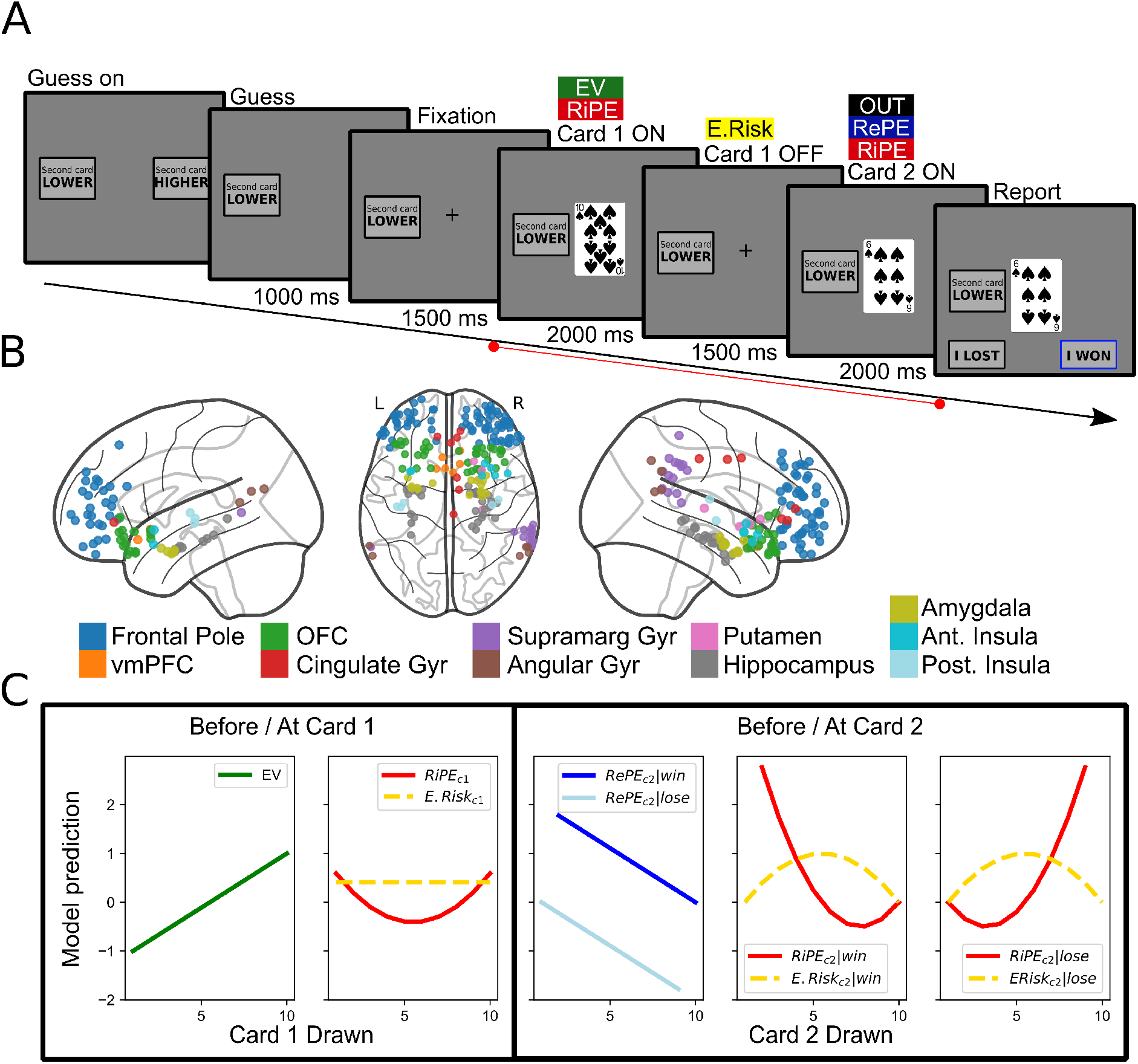
Experimental paradigm and computational model. (A) Overview of a single trial in the task completed by participants. Distinct trial events are labelled above, and each event duration is given below, the panels. The within-trial period used in subsequent neural analyses is denoted by the red line at the bottom, and a priori windows for each computational variable are overlaid in the coloured boxes. Neural analyses were conducted in epochs from -200 to 500ms around respective trial event onsets. (B) Contact locations across the pseudo-population. Each dot represents a bipolar sensor located at an interpolated coordinate between the paired contacts’ locations. Dots are coloured by their assignment to each atlas-defined region of interest (ROI). (C) Variation in computational variables as a function of the drawn number of card 1, conditioned on an initial guess that card 2 will be lower as illustrated in the example trial depicted in A. The left panel depicts variables predicted to arise before the onset of card 1 (EV: expected value, green; E.Risk; expected risk, yellow) and after the onset of card 1 (RiPE: risk prediction error, red). The right panel depicts variables predicted to arise before the onset of card 2 (E.Risk) and after the onset of card 2 (RePE: reward prediction error, blue; RiPE, all conditioned on card 2 actually being lower (i.e. a winning guess).

The task was designed in conjunction with a normative model of reward and risk [12, 4], allowing for the calculation of expected value (EV), outcome (OUT), expected risk (E.Risk), reward prediction error (RePE), and risk prediction error (RiPE) (Figure 1C). These variables are computed sequentially over the course of a single trial, based on information provided by the cards drawn (Figure 1A). After the second card is drawn, participants can compute RePE by comparing the actual outcome with EV and compute RiPE in terms of variance around E.Risk, thereby taking as input the RePE (see 4.8). Critically, each trial provides an independent sample of these variables, allowing us to employ single-trial decoding to unpack locally distributed multivariate representations

### 2.1 Behavior reflects knowledge of task structure and control for learning confounds

Our objective was to characterize the temporal properties of the neural representations underlying reward and risk-related computations. We aimed to study how multiple brain regions encoded these variables free from the potentially confounding effects of learning dynamics on neural activity. As such, we used a task that constrained the computational process within a single trial and did not elicit learning dynamics between trials. The task was designed such that the only optimal behavior was to report correctly at the end of each trial, and there was no optimal guessing strategy given that the deck was reset on each trial. We found evidence that participants indeed attended to the information provided by the presented cards as demonstrated by strong reporting accuracy significantly above chance in our sample (mean = 91.50 %, s.d. = 2.84 %, t(9) = 14.398, p = 1.184e-7; Figure 2A). We observed a trend towards decreased reporting accuracy on late trials of the task (Figure 2B) potentially reflecting effects of fatigue; however there was otherwise no effect of trial count on reporting accuracy (*β*_*t*_ = -1.272, z = -1.782, p = 0.075) and the accuracy remained high even in the late trials (mean = 88.4 %, s.e.m. =3.96 %). In all subsequent neural analyses we controlled for trials with inaccurate reports across the entire span of the task. Further, participants did not show any evidence that they adopted behavioral strategies commonly seen in learning tasks such as win-stay/lose-shift (*β*_*OUT*_ = -0.103, z = -1.719, p = 0.086; Figure 2C), nor did we find evidence that participants adopted other behavioral heuristics such as sticking to the same guess (*β*_*t* − 1_ = -0.092, z = 0.709, p = 0.478; *β*_*t* − 2_ = 0.005, z = 0.039, p = 0.969; Figure 2E) or left/right response key (*β*_*t* − 1_ = 0.181, z = 0.531, p = 0.595; *β*_*t* − 2_ = 0.166, z = 0.746, p = 0.456; Figure 2F).

**Figure 2:**
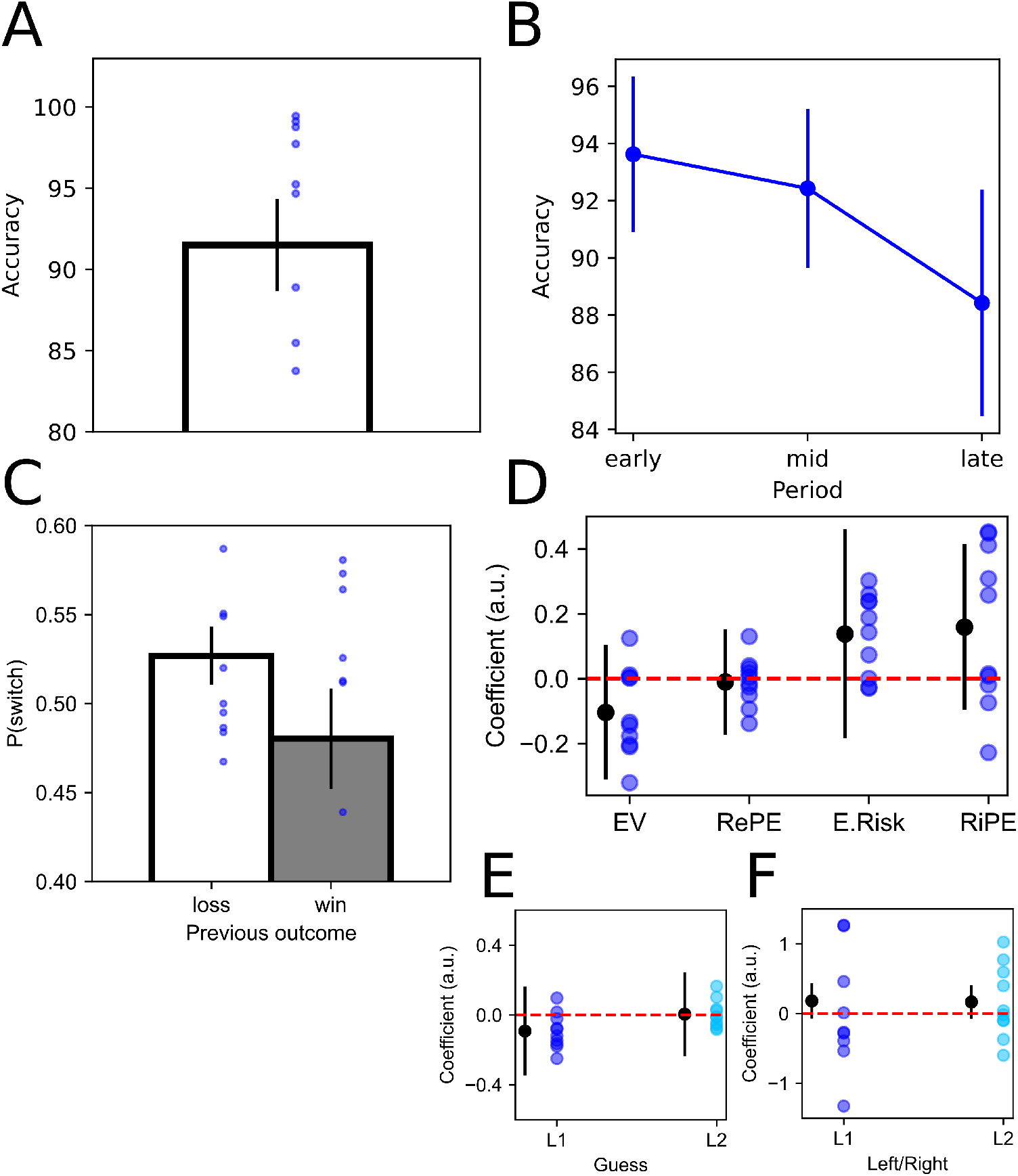
Behavior reflects understanding of task structure. (A) Overall average report accuracy across the task. Error bars depict standard error of the group mean (SEM) and dots are per-subject accuracy. (B) Average report accuracy over the span of the task. Early, mid, and late are defined by trial tertiles. Error bars depict SEM. (C) Proportion of trials in which participants switched their guess as a function of the outcome on the previous trial. Error bars depict SEM. No effects are significant, suggesting that participants understood the task and did not adapt any heuristic strategy (e.g. win-stay-lose-shift) in their guess. (D) Regression coefficients for the logistic model of predictors of guess choice. Black dots are overall fixed-effect coefficients and error bars depict 95 CI around coefficients. Colored dots depict per-subject random effects. Guess choice (E) and left vs. right response (F) auto-regressive model coefficients. L1 and L2 denote a lag of 1 and 2 trials back, respectively. Black bars denote fixed-effect coefficients and 95 CI, and colored dots are per-subject random effects. Along with D, no predictors are significant indicating that participants understood the structure of the task and were treating each trial independently.

Finally, given our interest in the underlying computations that emerge over the span of a trial, we tested whether computations on previous trials might have any influence on participants’ guesses on the following trial. We expected there to be no effect of the set of computational variables on the guess choice of the next trial if participants truly treated each trial independently. Critically, this analysis allowed us carefully test the contribution of reward and risk error on the next trial’s choice, where a lack of significant prediction would suggest that we succeeded in isolating the task from learning dynamics. Participants’ behavior was analyzed using logistic regression to assess whether guesses were predicted by reward and risk features of the previous trial, as index by the level of each of our variables of interest. We found that participants’ guesses were independent of their past experience both in terms of reward (*β*_*EV*_ = -0.104, z = -0.980, p = 0.327; *β*_*RePE*_ = -0.010, z = -0.122, p = 0.903) and risk (*β*_*E Risk*_ = 0.138, z = 0.842, p = 0.400; *β*_*RiPE*_ = 0.159, z = 1.225, p = 0.220; Figure 2D; see 4.8 for their definition). Again, no behavioral strategy was necessary to perform optimally given the task design; these results along with their high reporting accuracy provide evidence that participants understood the nature of the task well, and paid attention to both their guesses and the information presented through the span of a trial to report their outcome accurately. Importantly, this allowed us to proceed with confidence to the neural analyses which relied on the derived computational variables.

### 2.2 Decoding computational variables with multivariate neural signals

We aimed to coordinate the timing of reward and risk representations across multiple brain regions. Using a multivariate decoding approach, we examined the timing of each computational variable’s representation within a region of interest (ROI). Our design matrix for single-trial decoding allowed us to assess the decodability of each computational variable at its respective point of occurrence within a trial, leveraging the sequential structure of the task to temporally separate and de-correlate the variables. To ensure unique decoding of each variable of interest, we regressed out all other variables and covariates (e.g. per-trial report accuracy) from each neural feature. We then extended our decoding approach across ROIs spanning cortical and subcortical regions, focusing primarily in our key regions of interest but also conducting an exploratory investigation into a larger set of potentially relevant ROIs. By comparing the decoding results across regions, we aimed to identify temporal patterns of representation at a wide scale across the brain

### 2.3 Distinct modes of temporal organization between value-based computations

We hypothesized that the timing of decoding need not respect reward and risk domain boundaries, but could instead be organized in a manner consistent with the nature of the computation. Specifically, EV requires integrating over each remaining card’s reward likelihood after card 1 whereas OUT representations rely on exogenous information given when card 2 is shown, which in conjunction with the initial guess and card 1 determines trial success. One possibility is of sequential processing for outcome computations as the external information provided by card 2 is used in multiple ways to realize the trial’s outcome.

Consistent with a perspective highlighting differences in what the brain needs to do to compute each respective variable, we found different temporal configurations in the decoding patterns for EV and outcome across regions. We were able to decode EV in our hypothesized regions of vmPFC (p = 0.032) and amygdala (p = 0.022) in temporally overlapping windows (vmPFC [0.040 - 0.142 sec]; amygdala [0.044 - 0.102 sec]) after the onset of card 1 (Figure 3A,B). In an exploratory analysis, we were also able to decode EV in the hippocampus with two temporal clusters (early: p = 0.025; q = 0.117; late: p = 0.023; q = 0.117; Figure S1), with an earlier cluster overlapping in decoding timing [0.082 - 0.138 sec] when compared to vmPFC and amygdala.

**Figure 3:**
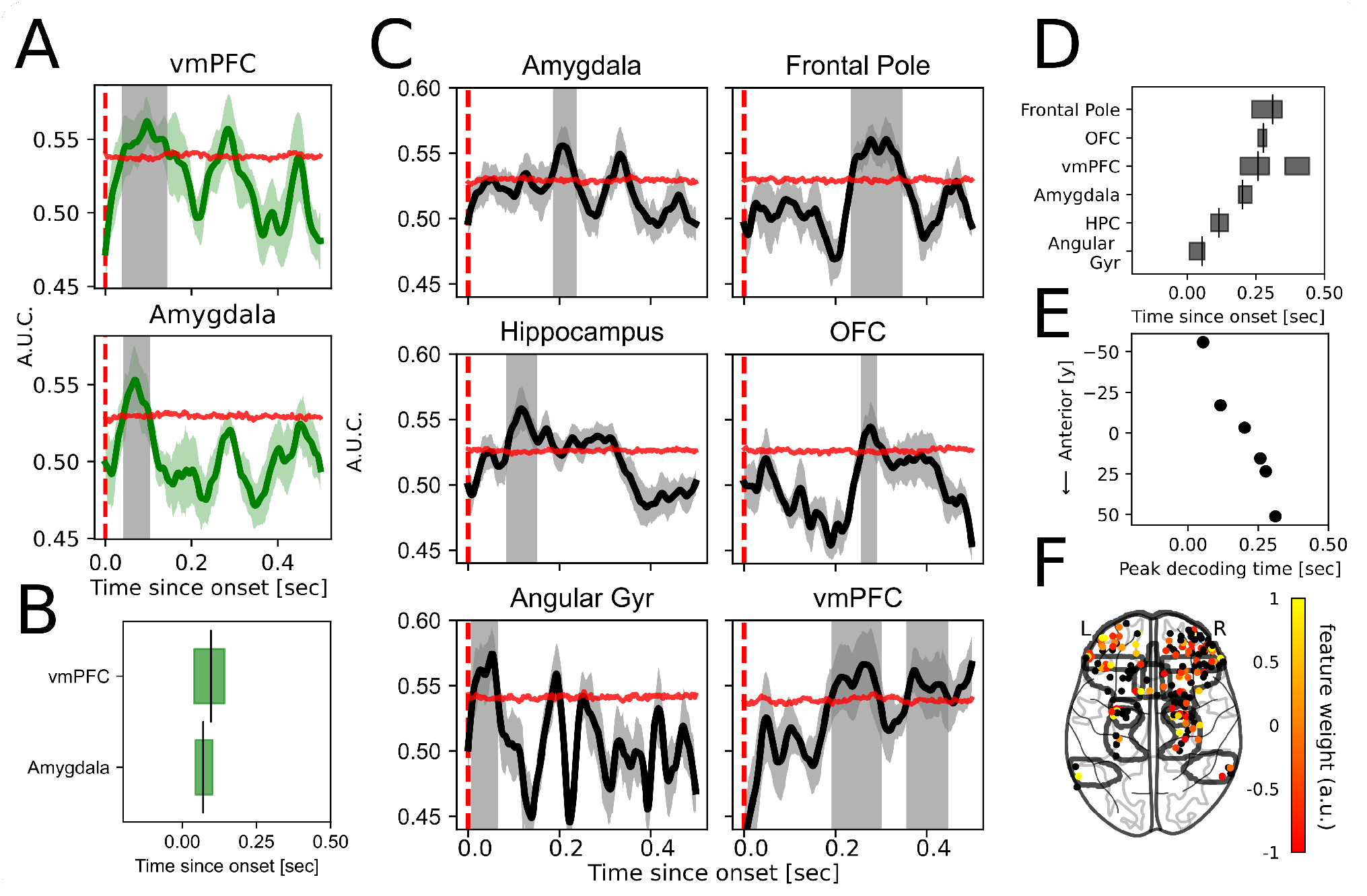
Parallel versus sequential temporal organization for value-based representations. (A) Decoding of expected value across contacts in each ROI, after the onset of card 1. Lines depict cross-validated receiver operating characteristic (ROC) accuracy (AUC), and shaded colored areas show standard error in AUC across folds. Horizontal red lines depict the 95th percentile of the permuted null distribution at each time point, and periods of statistical significance are shown in the shaded grey region (cluster corrected FWE < 0.05). (B) Timing of significant EV decoding for each ROI. Horizontal boxes depict periods of significant decoding and vertical bars indicate the time point of maximum significant ROC accuracy. All regions’ period(s) of decoding overlap, demonstrating a parallel temporal order. (C) In contrast, outcome decoding follows a sequential temporal organization across regions. Same as (A) but for outcome decoding in the period after card 2 is presented, when the participant has all information to know if their guess was correct and consequently if they won on that trial. (D) The timing of significant outcome decoding within each ROI follows a temporally cascading pattern. Boxes and bars are the same as (B). (E) Outcome decoding exhibits both temporal and spatial structure. Peak decoding times across ROIs follows a trend along a posterior to anterior axis as indexed by the y-plane centroid of contacts respective to each ROI. (F) Outcome is encoded in a locally distributed manner within each ROI. Each dot represents a bipolar source with its color depicting the feature weight, normalized across contacts within each ROI. ROIs for which outcome can be decoded in (C) are depicted by the black lines for reference.

Whereas EV decoding was significant in temporally overlapping extents across relevant ROIs, we observed a temporal cascade in periods of significant outcome decoding across regions of the brain (Figure 3D). We were able to decode outcome in a widespread set of regions across the brain, both in regions included in our *a priori* targeted ROIs, and those we surveyed in an exploratory analysis. Ordered from latest to earliest decoding time relative to the onset of card 2, we found significant outcome decoding in all prefrontal cortex ROIs including the frontal pole which incorporated anterior OFC (p < 0.001), more posterior lateral OFC (p = 0.02), and in two distinct temporal clusters in vmPFC / medial OFC (early: p = 0.011; late: p = 0.031). Consistent with our expectations based on prior work [32, 31] we were also able to decode the outcome in the amygdala (p = 0.020). In our set of exploratory ROIs we could decode outcome in both hippocampus (p = 0.002; q = 0.007) and angular gyrus (p = 0.002; q= 0.007). Consistent with the idea of a meso-scale level of spatial representation, we further found that within each ROI the multivariate contacts contributing positively to decoding the outcome were spatially distributed (Figure 3F).

Strikingly, we observed a spatiotemporal pattern in which the most posterior regions were decoded earliest, and this progressed sequentially to the most anterior region (Figure 3E). Here we characterized the anteroposterior location of each ROI by computing a projected centroid between the contacts respective to a given ROI, along the coronal (y) plane. We further tested whether there was a significant difference in the timing of decoding between pairs of ROIs, ordered by the relative latency of the peak of their respective decoding accuracy curves. For regions that exhibited two significant temporal clusters, we tested for differences in decoding latencies by using the most temporally proximal cluster with respect to pairwise ROIs. We found significant differences in timing between all pairs of temporally (and thus spatially) proximal ROIs, with outcome decoding first evident in angular gyrus [0.008 - 0.064 sec; y = -56], then hippocampus [0.086 - 0.150 sec; y = -17] (U = 957, p = 2.44e-18), then amygdala [0.188 - 0.236 sec; y = -3] (U = 825, p = 5.73e-17), then vmPFC [0.194 - 0.299 sec; y = 15] (U = 1108, p = 8.33e-7). While the period of outcome decoding in OFC [0.257 - 0.289 sec; y = 23] was significantly shifted later relative to vmPFC (U = 688.5, p = 8.14e-4), and earlier than frontal pole [0.236 - 0.345 sec, y = 51] (U = 620.5, p = 0.030), its temporal span fully overlapped with both neighbors despite their peak decoding latencies respecting this anterioposterior gradient.

Given the spatiotemporal structure of outcome representation across widespread areas of the brain, we investigated whether the directed connectivity between pairs of temporally adjacent ROIs differed as a function of the outcome on each trial, using a shuffle-corrected cross-correlation approach (see Supplementary Materials section 1.1). We hypothesized that areas earlier in the processing pathway as informed by the temporal ordering of outcome decoding would lead areas from which we could decode outcome relatively later. While we found evidence of differential cross-correlation between high and low outcome trials in prefrontal cortex, we instead observed that areas found to encode outcomes later led areas encoding outcomes earlier in time. Activity in frontal pole, which incorporated anterior OFC contacts, led those in the relatively more posterior lateral OFC (t(334) = -2.226, p = 0.027; Figure S5B), and activity in lateral OFC contacts led vmPFC activity in a similar outcome-dependent manner (t(24) = 2.165, p =0.041; Figure S5A).

### 2.4 Decoding of expected risk in OFC

We particularly hypothesized that we would be able to decode E.Risk in the anterior insula and prefrontal regions such as OFC given prior work implicating the role of these regions in uncertainty computations [4, 33, 34]. We also had strong temporal hypotheses of when in the trial E.Risk representations would be held: specifically in the period after the offset of card 1 (but before the onset of card 2) given that the computation of E.Risk takes as input the information from EV, which itself is computed once card 1 is shown. While we were able to decode E.Risk in OFC ([0.287 - 0.317 sec]; p = 0.028) in the predicted period after card 1 is offset, we did not find significant decoding of E.Risk in anterior insula (p = 0.397).

Furthermore, the temporal ordering of how multiple regions represent E.Risk remained equivocal. This is due in part to the lack of representation of E.Risk across multiple ROIs which would allow us to survey the widespread temporal organization of representation, though in our exploratory analysis we were able to decode expected risk in posterior insula ([0.102 - 0.154 sec], p = 0.022; q = 0.103) and in the supramarginal gyrus of the intraparietal lobule ([0.449 - 0.499 sec], p = 0.016; q = 0.103) (see Figure S1B). When considering the full set of ROIs in which we could decode E.Risk, we see a suggestion of sequential processing in the uncorrected significant cluster timings of our exploratory ROIs. Note too that an earlier sub-threshold increase in decoding accuracy in OFC (e.g. 0.080 - 0.166 sec in Figure 4) overlaps with the significant uncorrected significant cluster timing of posterior insula (Figure S1B, bottom panel)

**Figure 4:**
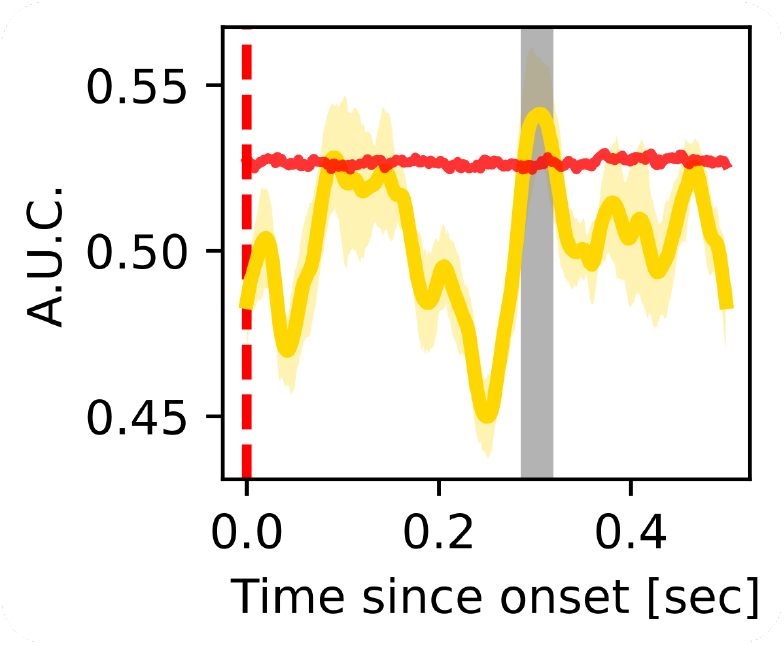
Expected risk can be decoded from the distributed contacts in orbitofrontal cortex. Lines depict CV ROC AUC, and shaded coloured areas show standard error in AUC across folds. Horizontal red lines depict the 95th percentile of the permuted null distribution at each time point, and periods of statistical significance are shown in the shaded grey region (cluster corrected FWE < 0.05).

### 2.5 Error computations across domains share a mixture of temporal configurations

We observed direct evidence of a mixture of parallel and sequential temporal ordering when we examined reward prediction error (RePE) and risk error (RiPE) representations. We observed significant decoding of RePE in OFC (p = 0.012), and anterior insula (p = 0.003) (Figure 5A), consistent with our hypotheses implicating both regions in representing this computational term. We observed that RePE representations extended to the posterior insula [0.269 - 0.299 sec] (Figure S2A), which we did not *a priori* predict to be implicated in this computation. Though the cluster in posterior insula did not survive correction (p = 0.025, q = 0.091), it overlapped in timing with the period of significant decoding in OFC [0.273 - 0.327 sec] and in the anterior insula an early partially overlapping cluster [0.277 - 0.307 sec] (relative to OFC: U = 160, p = 0.941; Figure 5C), which was followed by a later non-overlapping temporal cluster [0.339 20.405 sec] (relative to OFC: U = 952, p = 2.86e-18). Interestingly, in an exploratory analysis we found significant early decoding of RePE in cingulate gyrus [0.002 - 0.078 sec] (p < 0.001, q = 0.011; Figure S2A) which taken together with the decoding profiles in OFC, anterior insula, and posterior insula exhibit a mixed pattern of early decoding in one region followed by a multi-regional set of simultaneous RePE decoding (cingulate relative to OFC: U = 1092, p = 1.71e-19).

**Figure 5:**
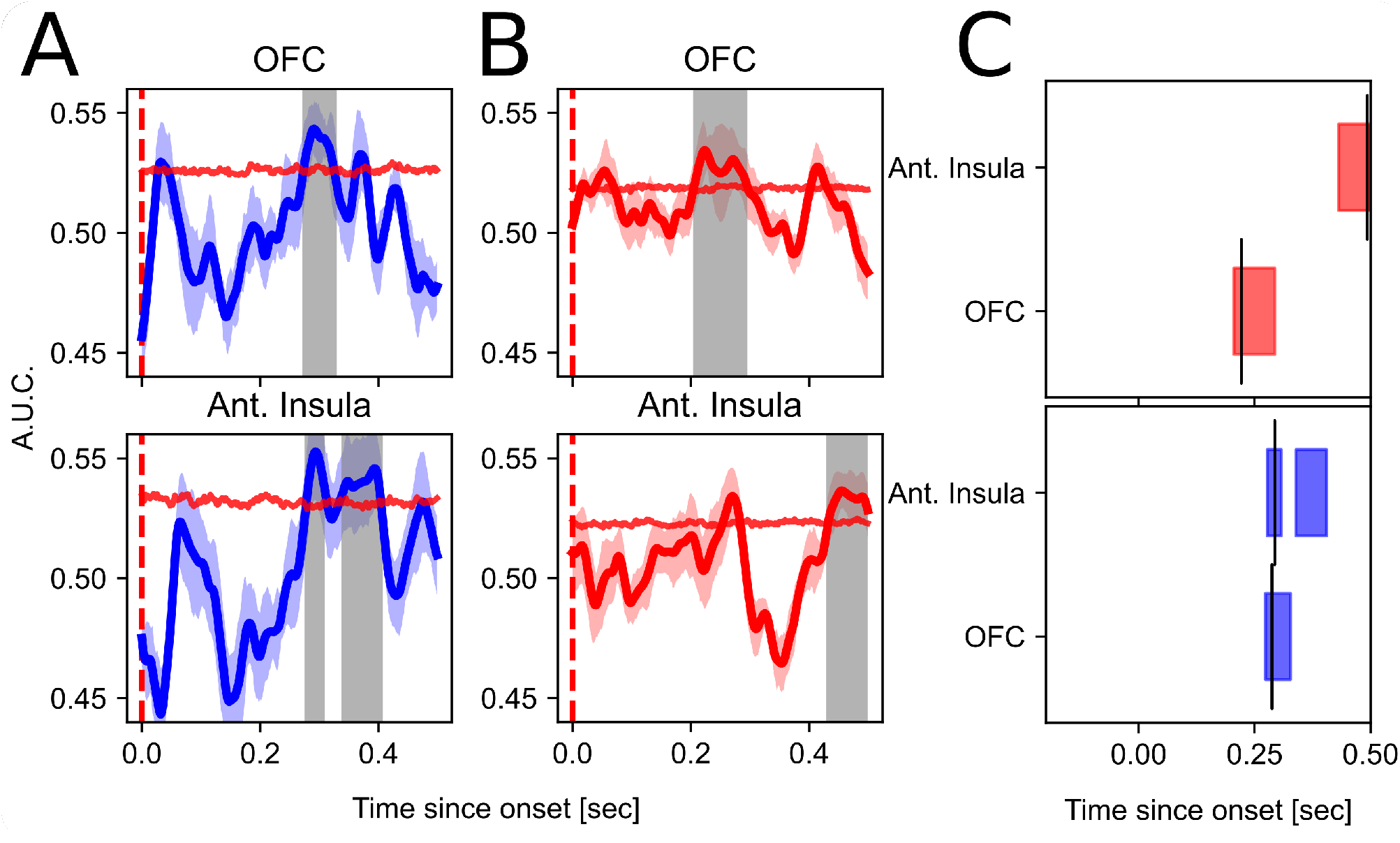
Decoding curves and periods of significance for reward prediction error (A; RePE) and risk prediction error (B; RiPE). In both variables across reward and risk domains we see a mixture of cascading and parallel decoding (see also Figure S2). (C) Timing of RePE and RiPE decoding periods across ROIs. RiPE decoding is shifted later with respect to RePE in anterior insula. Panel descriptions follow that described in Figure 3.

We held strong hypotheses that we would be able to decode risk error in the anterior insula, given prior work using the same experimental paradigm [4, 17]. Moreover, we hypothesized that the timing of decoding for risk error should occur later than that for RePE, given that in the computational formulation the risk error (RiPE) term takes as input the RePE (see 4.8. When we contrasted the temporal profile of RiPE decoding against that for RePE, we notably were able to decode RiPE in the anterior insula in a period shifted later in the epoch (Figure 5C), consistent with our predictions. Further, the temporal pattern of risk prediction error decoding across ROIs followed a similar temporal configuration characterized by early encoding in one region, here for RiPE in OFC [0.206 - 0.293 sec] (p < 0.001), followed by late decoding of RiPE across multiple regions including anterior insula [0.431 - 0.499 sec] (p = 0.016) (relative to OFC: U = 1620, p = 7.68e-24) and in an exploratory analysis the putamen [0.429 - 0.499 sec] (p = 0.024), with no differences in timing between the latter two regions (U = 647.5, p = 0.425).

Our analyses of error computations across reward and risk domains revealed a further temporal configuration, in which we able to decode multiple computational variables in overlapping periods within one region. In OFC, we were able to decode all computational variables designed experimentally to arise in the period after the onset of card 2: outcome (Figure 3C), RePE (Figure 5A), and RiPE (Figure 5B), with overlapping temporal extents [0.273 - 0.289 sec].

### 2.6 Reward error representation generalizes to risk error in Anterior Insula but not OFC

Among regions of theoretical interest, the OFC and anterior insula would stand out to be of particular relevance for representation multiple computations given their established involvement in both reward and risk processes (e.g., [33, 4, 42]). Consistent with this prediction, we were able to decode both reward error (RePE) and risk error (RiPE) in both anterior insula and OFC, despite the two regions being distinguished by the sequential versus simultaneous decoding of these variables, respectively. Highlighting again the conceptual importance of characterizing the temporal organization of neural representation, here we directly tested for evidence that neural activity in OFC and anterior insula could hold a representation of one input variable for the purpose of a second downstream computation; specifically, whether the representation of RePE might be relevant for decoding RiPE.

We ran a generalized decoding analysis to directly test for the possibility that the neural representation of one input variable (e.g. RePE) can be leveraged for the purpose of computing a downstream variable (e.g. RiPE). We first focused specifically on signals in the anterior insula given prior work emphasizing its role in risk error computations [4, 17, 37], and the evidence that we were able to decode both variables individually in the anterior insula, in which the timing of RePE decoding preceded risk error (Figure 5). Consistent with the idea that the brain needs to first compute the reward prediction error before using this information to compute the risk error, we hypothesized that activity in the time window of RePE representation would be able to predict activity related to RiPE at a relatively later period in the same epoch. We constrained our decoding analysis here to the period in the trial relevant to RePE and RiPE representation (0.200 20.500 sec after the onset of card 2) and tested for generalization both across variables (i.e. trained on reward error and tested on risk error) and time. Indeed, we found evidence of significant cross-validated generalized decoding by which a model trained on anterior insula activity signal in a relatively earlier period [0.200 - 0.391 sec] was able to predict risk error later in the epoch [0.383 - 0.499 sec] (p = 0.014; red cluster in Figure 6A). Indeed, the temporal extent of significant cross-variable training and testing overlapped with the decoding periods for RePE and RiPE, respectively, as described in 2.5. We continued to find evidence of significant generalization when tested against a more stringent statistical threshold (trained on RePE: [0.200 - 0.283 sec]; test: [0.439 - 0.489 sec]; p < 0.001, orange cluster in Figure 6A), and when we did not constrain the time window for analysis (see Figure S3).

**Figure 6:**
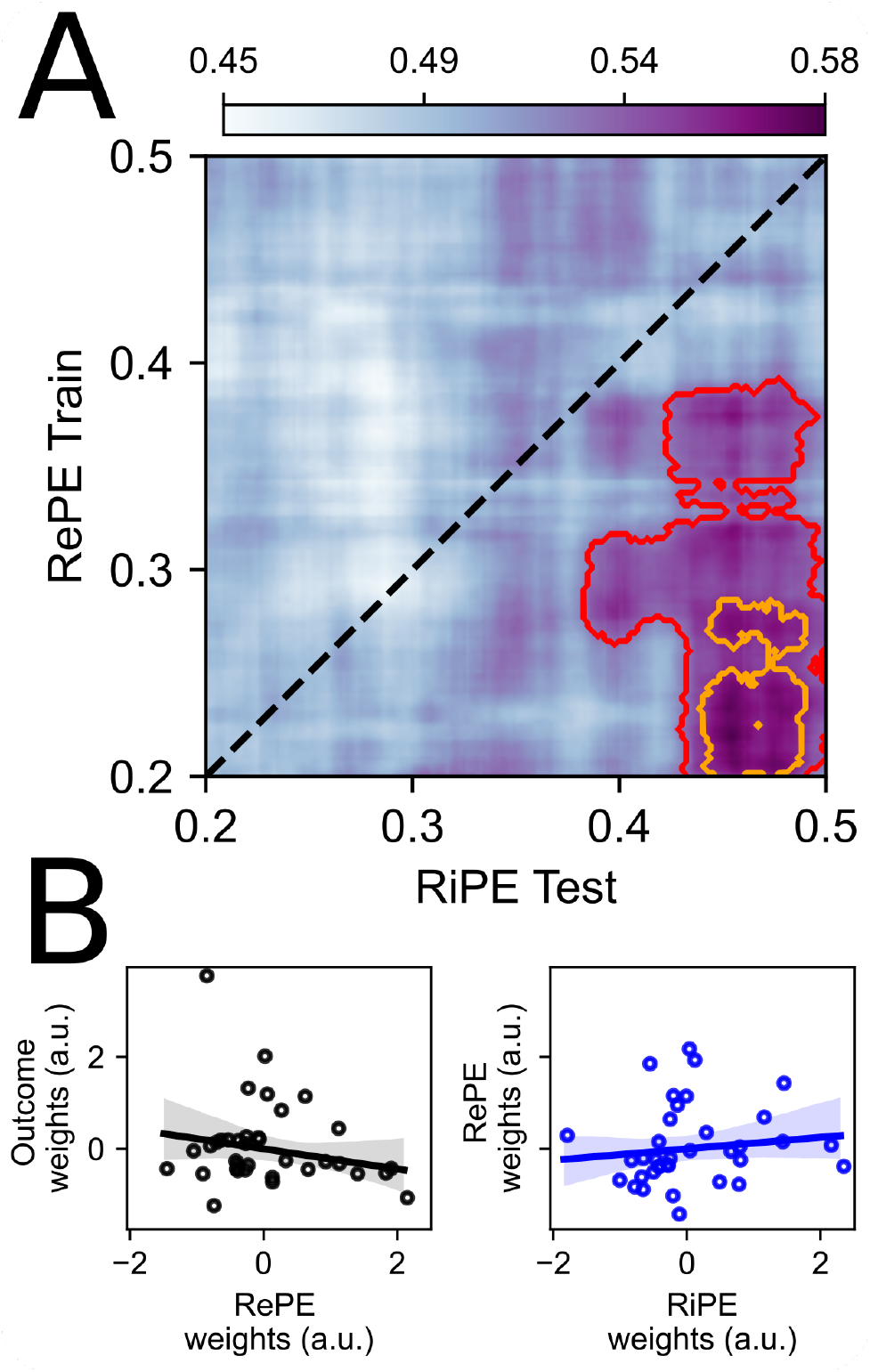
Generalizing and independent representational roles in anterior insula and OFC. (A) A model trained on multivariate signal in anterior insula encoding RePE early in the epoch after the presentation of card 2 can generalize across time and computations to decode RiPE later in the same epoch. Significant decoding areas are highlighted by the red (p = 0.01) and orange (p = 0.002) contours (both cluster corrected FWE < 0.05). (B). OFC feature weights were not correlated across variables, both those respective to the decoding of outcome and RePE (r = -0.206, p = 0.228) and to the decoding of RePE and RiPE (r = 0.126, p = 0.463). Axes denote the normalized feature weights averaged over the same period for all variables [0.273 - 0.289 sec after the onset of card 2].

Our predictions regarding the contribution of simultaneous decoding of multiple variables in OFC was less concrete. On one hand this parallel temporal configuration could reflect an interaction between RePE and RiPE within the same distributed population, in which case the neural representation for one variable would contribute to decoding another variable. Alternatively, our finding of simultaneous decoding across variables could reflect independent encoding of these variables along different distributed codes within the same OFC ROI. Contrary to the first potential mechanism, when we conducted the same generalized decoding analysis we did not find evidence of cross-variable decoding at any time point in OFC, both when we trained on outcome and tested on RePE (Figure S4A) and when we trained on RePE and tested on RiPE (Figure S4B). We proceeded to directly test the possibility that independently distributed codes within the OFC were implicated in the representation of each computational variable. For a given variable, we characterized the contribution of each OFC contact source to the significant cross-validated decoding scores reported above using a leave-one-feature-out approach for measuring feature weights, averaging over the period of overlap between outcome, RePE, and RiPE [0.273 - 0.289 sec]. We then tested whether the representational vectors capturing the distributed code in OFC correlated across variables. We found no significant correlation between features weights that encoded outcome and RePE (r = -0.206, p = 0.228), nor between RePE and RiPE (r = 0.126, p = 0.463), which together with our decoding generalization results supports the notion of independent representations of outcome, RePE, and RiPE in OFC (Figure 6B).

## 3 Discussion

We set out to investigate how the human brain represents a host of computational variables relevant for decisionmaking in both value and uncertainty domains. Specifically, we aimed to elucidate the temporal properties of neural representations supporting computations of reward and risk expectations and errors. We were able to decode all our computational variables of interest in wide-spread regions. Critically, we distinguished two modes of temporal organization by which computations are represented in the brain. In the reward domain, expected value and reward outcome differed in their temporal profiles with the former encoded in parallel within a similar time frame across regions, while outcome representations followed a sequential spatiotemporal cascade from posterior to anterior brain regions. Thus, while expected value and outcome are encoded in overlapping distributed value-based networks [13, 43, 44], these two distinct value-related variables exhibit very different spatiotemporal properties. Moreover, error computations for both reward and risk shared a similar temporal profile reflecting a mixture of sequential and parallel encoding. We demonstrate the importance of elucidating the timing of different types of computational representations by presenting evidence that the distributed code in anterior insula relevant for reward error precedes and directly contributes to decoding risk error. In contrast, representations related to outcome, reward prediction error, and risk error are distinct in OFC. Together, our findings highlight distinctive patterns of timing by which the brain represents reward and risk variables, as well as potential functional roles of neural representation in regions such as the anterior insula and OFC.

We recorded human neural activity across the brain using intracranial electroencephalography, which is well situated to describe an intermediate level of analysis with broad spatial coverage across the brain [45, 41] and relatively high specificity within a region [38]. We took an multivariate decoding approach in which we examined whether the distributed activity across contacts within a specific ROI was able to decode our computational variables of interest, and we applied this approach across multiple regions. We were motivated theoretically by existing work on reward and risk encoding in human iEEG [33, 39, 34], which had taken an encoding approach and focused predominantly single or a small subset of regions. We extended this work not only by exploring a larger set of regions in our exploratory analyses, but importantly by elucidating the temporal organization of reward and risk representation across the brain.

Critically, we leveraged an experimental design in conjunction with a normative model of reward and risk which separated, in time over the span of multiple sequential trial events, when the information relevant for each computational processes was available. We could therefore examine specific periods of neural data with respect to each computational variable, in the span of a single trial, allowing us to look at the the unique decodability of a set of computational variables which are not often de-coupled in other experimental contexts. By regressing out variance attributable to other computational variables in each decoding scheme, we further ensured that the neural data going into each decoding model did not carry residual information correlating with variables other than the one of interest to be decoded. Moreover, our task structure allowed us to investigate the degree to which neural activity could decode our computational variables at the level of a single trial [46] with high temporal specificity, and allowed us to control for the effect of learning dynamics on neural representations. We could therefore examine isolated, learning-independent computational features.

Our findings of both parallel and sequential modes of temporal organization are consistent with experimental work in other domains of cognitive neuroscience generally [18, 20] and in the domain of value-based decision-making specifically [15]. We found a unique temporal cascade in the regions that represented the outcome of each trial in a manner that respected an anteroposterior gradient. Signal in the angular gyrus ROI in the posterior parietal cortex could be used to decode outcome first; this then proceeded sequentially across ROIs to the frontal pole over the period in which all information for knowing the outcome was available. While previous work has reported outcome correlates over widespread regions of the brain [47, 48, 43, 49], here we exploited the high temporal resolution of human iEEG to show that outcome representations across the brain have a distinct spatiotemporal profile. The gradient-like organization from posterior to anterior regions of the brain is consistent with similar architectures reported across multiple regions for diverse cognitive processes [50, 51, 52]; here we show that this topographical arrangement is widespread (from parietal to prefrontal regions) and temporally ordered in the representation of reward outcome. In the context of this task, the outcome (i.e., the accuracy of the initial guess) is unique in that explicit feedback is not presented to the participants. Instead representing the outcome entails processing information conditioned on the guess, card 1, and finally card 2. Exploiting multiple pieces of observable information to derive the resulting outcome could potentially drive this widespread cascade in encoding. Our further finding of anterior to posterior outcome-related directed connectivity between subregions of the PFC might be reflective of resulting top-down processing [53], in which existing expectations of reward and risk can be computationally resolved once the outcome is known.

Conversely, regions in which we were able to decode expected value (EV) all showed overlapping temporal spans of significant decoding, after the onset of the first card when enough information was available for participants to form a conditional expectation of the probability of guessing correctly on that trial. Spatially, our findings of EV decoding were consistent with previous work that reported representations of EV in vmPFC [39], amygdala [54, 29, 32], and hippocampus [55, 56]. Here we extend this previous work further not only by showing that EV is coded in the distributed, multivariate activity within each region, but also showing that the code is held in a parallel temporal configuration. The EV term as defined in this context requires holding multiple pieces of information in the form of the reward probabilities associated with each possible second card, and the internal process of integrating over this information. One possible interpretation of simultaneous EV representation across regions is that this reflects the top-down nature of EV computation in which the relevant information (i.e., each possible draw for the second card and their respective likelihood of reward) does not require further external input but relies instead on held knowledge of the instructed task set. Interestingly, this temporal configuration has been discussed as a potentially important neural organization across cognitive domains [57] and scales of analysis [58, 59, 60].

We also sought to investigate the potential functional advantages conferred by temporally organized computational representations in the brain. One possibility is that multiple regions might all hold a similar piece of computational information (e.g. reward prediction error [RePE]), but that different regions use this same information for different purposes. For example, our ability to decode RePE from activity in cingulate cortex rapidly after the onset of card 2 is consistent with the literature on error signals in dorsal anterior cingulate cortex (dACC), both measured with intracranial [61] and scalp [62] EEG. This fast temporal signature, in conjunction with prior work on cingulate encoding of error signals in other domains [61], suggests that fast decoding of RePE we find in cingulate cortex might reflect a more general error (in the form of deviation from expectation).

In contrast, we show that the encoding of RePE in anterior insula serves a specific functional role that directly relates to later RiPE computations in the same region. We show a relationship between the neural code supporting RePE and RiPE, both indirectly via the shifted temporal profile between RePE and RiPE decoding in anterior insula over the same period, and directly with our generalized decoding analysis in which the neural representation of reward error at an earlier period can significantly decode risk error later in the trial. This finding speaks to the notion of neural representations progressively ‘building’ computations, consistent with a line of evidence emphasizing the role of specific neural activity in composing computations from their constituent components [47, 63, 48]. Here we extend beyond established associations between anterior insula and risk error [4, 37] and show that the distributed activity in anterior insula plays a further crucial role in leveraging input computations (i.e., RePE) to decode risk error. In other words, we adopt the notion of neural representations of computations being compositional, and present evidence of this across the value and uncertainty domains. This nonetheless does not preclude the possibility of independent representations of multiple types of computations in non-overlapping distributed activity within a region, as we show in independent representations of outcome, RePE, and RiPE in OFC.

Further highlighting the importance of resolving computational timing at a fast timescale, we were able to decode expected risk decoded later in the trial prior to the onset of the second card consistent with the description of expected reward and risk in our normative model. Our results were also congruent with previous reports in human [34] and animal [63, 64, 65] electrophysiology studies on the relative timings between expected reward and risk encoding, in which expected reward is encoded shortly after cue presentation whereas expected risk encoding is evident in activity starting before the period in which the outcome is known. However, we found a posterior versus anterior insula distinction in the decoding of expected risk and risk error, respectively, in contrast to a previous fMRI study using the same task [4] which reported spatial distinctions between expected risk and risk error correlates all within the anterior insula. One possible explanation for this discrepancy is that our recording contacts lie along the inferior plane of the anterior insula, more consistent with the reported BOLD correlates of risk prediction error than to that of expected risk. Another possibility speaks to potential discrepancies in the computational information encoded in human electrophysiology versus fMRI, an avenue of research to be explore more thoroughly in future studies.

Here we provide one picture by which the brain encodes multiple features relevant for value-based decisions, in which neural representations of computational variables are temporally organized and spatially distributed. Our findings also contribute to an emerging literature describing how neural implementation of different reward and risk computations unfold in time interactively. Importantly, while our normative model afforded us the ability to look at computational representations isolated from learning dynamics, further work extending our approach to subjectively computed computational variables, for example through learning or belief updating mechanisms, will enrichen our understanding of how the human brain represents the many facets of information important for complex decisions in the world.

## 4 Method

### 4.1 Participants

Our sample included patients-subjects (n=10, 7 female, age range 22-56, mean = 37.70, s.d. = 9.93) who underwent neurosurgical implantation of intracranial depth and surface electrodes to enable chronic evaluation and localization of seizure foci for the treatment of refractory epilepsy. Within our full sample, 6 subjects were implanted only on the right side, 3 were implanted on the left, and 1 was implanted bilaterally. All data were collected in periods free from seizure actiivty at least 1 hr before and after the experiment and data were visually inspected for interictal activity characterized by stereotyped transients in the raw recording. All research protocols were approved by the University of Iowa Institutional Review Board. Subjects provided written consent prior to participation in research and could rescind consent at any point without consequence to their medical evaluation.

### 4.2 Experimental paradigm

Patient-subjects were presented with two cards drawn sequentially and without replacement on each trial. The cards were shuffled from a deck of cards comprised 10 cards (ace to ten) and excluding face cards. Participants were instructed to treat an ace card as denoting “1”. The cards were reshuffled after every trial. Prior to the drawing of the first card, participants were instructed to predict whether the second card would have a higher or lower numerical value than the first.

Participants were given up to 7 seconds to respond. Their guess was displayed on the screen for a duration of 1000 ms, and a pre-card fixation cross was shown for 1500 ms. We then drew the first card which was displayed for 2000 ms before a 1500 ms inter-stimulus period in which only their intial guess remained on the screen. The second card was then shown for another 2000 ms. Upon the presentation of the second card, all information necessary to determine the accuracy of the initial guess was available to the participant. Finally, participants were prompted to report whether they had won (guessed correctly) or lost (incorrectly). Note that the initial guess of the participant remained on the screen throughout the presentation of both cards as well as the report period to reduce memory demands. A schematic of the trial structure and event timings are presented in Figure 1A. To incentivize participants to respond both to the initial guess and the final report, participants were instructed that they would receive a reward outcome in the form of 10 game points if their guess was correct and a deduction of 10 points from their cumulative total if their guess was incorrect. Correct or incorrect reporting of their guess accuracy at the end of the trial further resulted in gain or loss of 5 points, respectively. While these outcomes were displayed during the initial instructions and training block, during the task period in which data were analysed these outcomes were not presented to the participant.

### 4.3 Procedure

Participants remained in the epilepsy monitoring unit of the Epilepsy Center at the University of Iowa Hospitals and Clinics for 2-4 weeks after implantation, under clinical direction. The experiment reported here was delivered during this monitoring period. The epilepsy monitoring unit included a recording facility in an electromagnetically shielded room. Participants were awake and in a hospital bed or arm chair for the duration of the experiment, including during the delivery of task instructions. Participants were simultaneously provided verbal instructions and written instructions programmed into the task code which comprised static messages describing what the participants needed to do during the task (see 4.2) as well as interactive exercises to reinforce key task details, familiarise participants with the behavioural demands of the task (i.e. guesses and accuracy reports), and elucidate the incentive structure of a trial. Participants responded using left/right keyboard presses for both the guess and report; left/right mappings were randomised on every trial. To motivate participants to attain the most points they could, participants were instructed that they competed against previous players to rank on a leaderboard based on the total accumulated points at the end of the task. After participants were instructed and verbally confirmed they understood the task, they proceeded to complete 1 (n=4) or 2 (n=6) sessions comprising 90 trials per session; the number of completed sessions was based on participant willingness. We thus recorded a total of 16 sessions and 1440 trials across the sample. A subset of the participants (n=2) had previous exposure to instructions and experience with the task as they completed pre-implantation function MRI scans.

### 4.4 Recording

We used a combination of depth electrodes comprising low impedance clinical contacts (2.2 mm - 10 mm intervals), acquired using the Neuralynx (Bozeman, MT) Atlas system, and simultaneous multi-contact subdural grid electrodes embedded in a silicon membrane (Ad-Tech Medical Instrument, Racine, WI). The electrophysiological data were recorded at a sampling rate of 2 kHz with 24-bit resolution and bandpass filtered during acquisition between 0.1 to 500 Hz.

### 4.5 Imaging and contact localisation

Before and after electrode implantation, we collected high-resolution structural MRI data of the brain for each participant. For the first subset of participants (n=5), pre-implantation structural images were acquired from a 3T GE Discovery MR750w scanner with a 32-channel head coil (T1: FSPGR BRAVO 1.0 mm^2^, slice thickness = 0.8 mm, FOV = 256 mm^2^, TR = 8504ms, TE = 3.288ms, FA = 12^*°*^, TI = 450). The second subset (n=4) completed pre-implantation structural scans on a 3T GE SIGNA Premier scanner with a 32-channel head coil (T1: FSPGR BRAVO 0.8 mm^2^, slice thickness = 0.8 mm, FOV = 256 mm^2^, TR = 8576ms, TE = 3.364ms, FA = 12^*°*^, TI = 900). One participant completed their pre-implantation scan on a 1.5T Siemens Avanto with a 32-channel head coil (T1: MPRAGE 1.0 mm^2^, slice thickness = 1.5 mm, FOV: = 256 mm^2^, TR = 2200ms, TE = 2.96ms, FA = 8^*°*^, TI = 450). Patients completed a post-implantation T1 scan on a 3T Siemens Skyra with a T/R head coil (MPRAGE 0.98 mm^2^, slice thickness = 1mm, FOV = 256 mm^2^, TR = 1900 ms, TE = 3.44 ms, FA = 10^*°*^). Preoperative structural MRI images were co-registered to post-implantation structural MRI images, guided by post-implantation computed tomography (CT) scans (in-plane resolution 0.5 × 0.5 mm, slice thickness 1-3 mm), using custom MATLAB (Mathworks, Natick, MA) scripts and affine registration (FSL’s FLIRT, Jenkinson et al., 2012), and all images were processed to 1mm^3^ isotropic resolution. Visual comparison with intra-operative photographs was conducted to verify accuracy. For contact coordinate localisation, each participant’s brain was co-registered to the CIT168toMNI template brain [66] using ANTs software [67], and resulting contact locations are shown against the template (see Figure S8A).

### 4.6 Preprocessing

Intracranial electroencephalography (iEEG) recordings were preprocessed using custom scripts relying on functions from the MNE-Python toolbox [68]. The data were trimmed to include only periods in which participants were doing the task, downsampled to a sampling rate of 500 Hz (i.e. 1 sample every 2 ms), and line noise was removed using a 60 Hz notch finite impulse response filter. The data were high pass filtered at 1 Hz, entered into an independent components analysis (ICA [69]) denoising step, and then low-pass filtered at 250 Hz. The ICA-based denoising approach involved computing an ICA on the continuous contact timeseries, extracting the absolute value weight matrix from the ICA decomposition, and comparing the distribution of weights over contacts against an uniform distribution using the Kullback-Leibler divergence (*D*_*kl*_; [70] given evidence that systematic sources of noise and volume conduction effects can affect multiple contacts and to isolate spatially specific sources [71, 72, 73]. Independent components corresponding to weight distributions reflecting uniform distributions, defined with respect to a *D*_*k l*_ threshold, were visually inspected and removed from the data. This was done at two-stages: across all contacts on all electrodes within-subject (*D*_*kl*_ threshold = 0.2) to detect global sources of noise, and across all contacts along each electrode (*D*_*kl*_ threshold = 0.05) to detect local sources of noise permeating across an electrode. Finally, contacts sharing an electrode were re-referenced along bipolar pairs, identified by adjacent contacts from the same electrode, to further isolate local activity around the contact locations and mitigate the effects of volume conduction [74, 71]. Bipolar pairs were excluded if both contacts were in white matter as defined by a segmentation of the CIT168 template using FSL’s FAST algorithm [75]. The template-space coordinates between bipolar pairs was interpolated for visualization (see Figure1B and Figure S8B). The preprocessed data were epoched at within-trial events: the guess response, the onset of card 1, the offset of card 1, the onset of card 2, and the report response (1A. Epochs for all events were defined by a period spanning -200ms to +500ms around the event onset, with the period before event onset defined for baseline correction. The epoched bipolar timeseries were entered into all subsequent neural analyses.

### 4.7 Behavioural analysis

To survey the degree to which participants paid attention and completed the task, we analyzed behavior using mixed-effects regression models to differentiate between within- and between-subject sources of variance, specifying random intercepts and slopes for all terms, for each participant (shown in Wilkinson-Rogers notation). First, we tested whether their reported accuracy *R* different significantly from chance (50% accuracy), with accurate responses in the behavioral report data defined as correctly selecting whether they won or lost on a trial:

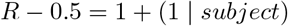

Using a mixed-effects logistic regression model, we tested whether report accuracy tended to decrease over the span of the task by regressing the report accuracy onto the trial number (scaled between 0 and 1) within a block. Here and below proportionality represents the log-odds (i.e. logit) function on the dependent variable:

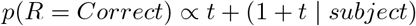

To examine whether participants were influenced on each trial by previous computational quantities, we modeled the guess choice on a trial (*G*; whether the second card would be higher or lower) as a function of the computational variables of interest on the previous trial: expected value (EV), reward prediction error (RePE), expected risk (E.Risk), and risk error (RiPE):

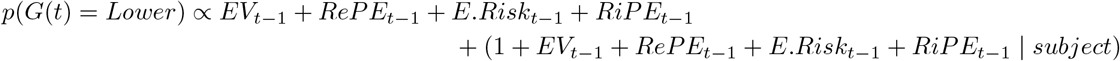

We assessed whether participants’ choices were influenced by previous events (e.g. win-stay/lose-shift across high/low guesses). To do so, we re-parameterized their choice on a given trial in terms of switch (different from the guess on the previous trial) or stay, and regressed switch/stay responses onto the outcome of the previous trial:

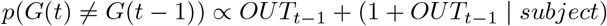

To further test whether participants understood the independent nature of each trial, we constructed models examining guess choice and left/right key press (*K*) stickiness, extending two trials back:

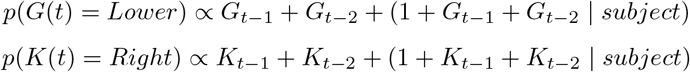

### 4.8 Computational analysis

Before the onset of the first card, the prediction about the number on card 1 (and thus the probability of reward given the guess) does not vary with the initial guess, so is denoted following [12] as a constant *P*_0_ and excluded from subsequent analyses. Upon the onset of the first card, participants are able to compute the *EV* : the expectation of the probability of reward conditioned on the value of card 1 and their guess

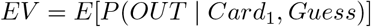

This term can be computed in a normative manner given the relatively limited set size of 10 total values and the knowledge that cards are drawn without replacement, meaning that the second card cannot be the same as the first. For example, if the participant guessed that the second card will be higher and drew a 9 on the first card, it is intuitive that the probability of getting a reward (i.e. that their guess will be correct) will be low given that 8*/*10 of the remaining cards available to draw as the second card are lower than the first. The computation of this *EV* term also allows for an error term at this stage, defined as the deviation between the *EV* (the expectation about the probability of reward given the guess), and the constant *P*_0_ (the probability of reward given the guess). In practice this error term is perfectly correlated with the *EV* term since *P*_0_ is a constant, and is not included in subsequent analyses.

Note that prior to the onset of card 1 we can formalise an expected risk term. Before participants see the first card, they can already form an expectation about the risk around the eventual expected value by integrating over the squared difference between all possible *EV* s across the 10 values available for card 1 and the constant *P*_0_:

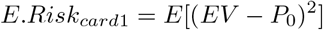

However, as with *P*_0_, *E*.*Risk*_*card*1_ does not vary across trials since it is not conditioned on the value of card 1. Nevertheless, it is useful for illustrating the risk error, a higher-order uncertainty defined as the difference between the square deviation from expected value (i.e. risk or unsigned prediction error [10]) and expected risk at the onset of card 1:

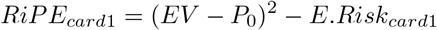

We designed the task to include a brief period of 1.50 sec in which the first card is offset from the screen but before the second card is shown (Figure 1A), with the assumption that expectations about the risk around card 2 would be computed in this period [12]. Here we similarly define *E*.*Risk*_*card*2_ in terms of the expectation about the risk around the expected value, where risk is again defined as the eventual square deviation of the outcome *OUT* from expected value (i.e. reward prediction error):

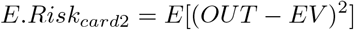

Finally, when the second card is presented the participant is able to computationally resolve their previous estimates of the probability of reward as the combination of their guess, the value of card 1, and now the value of card 2 provides all necessary information to know the outcome, and thus the computation of reward prediction error:

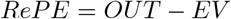

The RePE accordingly informs the risk prediction error term at card 2:

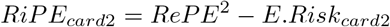

### 4.9 Region of interest definition

We specified a set of 5 *a priori* regions of interest (ROI) spanning frontal and subcortical regions given our strong hypotheses of the role of the prefrontal cortex (PFC), insula, and amygdala in value and uncertainty based computations, using the Harvard-Oxford (H-O) probabilistic atlas to delineate contacts respective to each ROI. The 5 *a priori* ROIs included 3 sub-regions of the PFC: Frontal Pole, lateral OFC (Frontal Orbital Cortex), and vmPFC (Subcallosal Cortex) where parentheses refer to H-O labels. Note that the “Frontal Pole” ROI label as defined by the H-O atlas included contacts in anterior lateral OFC (Supplementary table 1) whereas the “Frontal Orbital Cortex” label included contacts in posterior lateral OFC (Supplementary table 3), and the “Subcallosal Cortex” label included contacts in medial OFC (Supplementary table 2). We also hypothesized that the anterior area of the insula (Insular Cortex) would be particularly relevant for risk representations given findings from prior work [4] using the same experimental paradigm. Since the H-O anatomical mask for the Insular Cortex comprised voxels spanning both anterior and posterior aspects of the insula, we split the anatomical mask at MNI(2mm) Y = 63 ensuring that all contacts ascribed to anterior and posterior were respectively located on positive and negative coordinates in template space along the Y plane (See Supplementary Tables5 and 6).

We were also interested in exploring whether reward and risk representations could be held in wide-spread regions across the brain outside of our main hypothesized ROIs. We thus included an additional set of 6 ROIs comprising: Posterior Insula, Putamen, Cingulate gyrus, Hippocampus, Angular Gyrus, and Supramarginal Gyrus. The contact locations within each labelled ROI and their corresponding Brodmann area labels are presented in the supplementary tables (see Supplementary Materials section 1.2). ROIs were defined anatomically according to the H-O atlas as implemented in FSL and mask boundaries were defined at a voxel threshold of 10 (p = 0.1). Contacts within the boundaries of the thresholded probabilistic maps of > 1 ROI were assigned according to the max probability, and contacts across bilateral ROIs were aggregated.

### 4.10 Decoding analysis

#### 4.10.1 Feature preprocessing

We set up an encoding model to preprocess each of our features (contacts) for the decoding model. At the within-subject level before concatenation across subjets (see 4.10.2), for each feature we defined a general linear model (GLM) to regresss out covariates of no interest, as well as computational variables aside from the current variable of interest for decoding. The design matrix of the GLM included indicator vectors for the onset of the cards, the offset of card 1, and mean-centered parametric vectors for respective (i.e., not currently decoded) computational variables: *EV*_*card*1_, *OUT*_*card*2_, *ReP E*_*card*2_, *E*.*Risk*_*card*1*Off*_, and *RiP E* along with the observed risk *O*.*Risk* (i.e. squared *ReP E*), aligned to the trial event denoted in the subscripts. *RiP E* is computed at both card 1 and card 2 onsets (see 4.8); quantities at each of these trial events were accordingly aligned in the design matrix. The per-trial report accuracy was included as a covariate, aligned at the report response event. Because of colinearity between *Out*_*card*2_ and *ReP E*_*card*2_, and between *O*.*Risk* and *RiP E* (see Figure S6B), we orthogonalized the latter variable with respect to the former for each correlated pair such that the residual variance in *ReP E*_*card*2_ and *RiP E* were unrelated to *OUT*_*card*2_ and *O*.*Risk*, respectively. We subsequently dropped the variable of interest, regressed each feature vector against this covariate GLM, and replaced each feature by its respective residual vector.

#### 4.10.2 Pseudo-population design

For each ROI and participant, we set up an expanded pseudo-population design to create the feature matrix for decoding. First, we concatenated the epoched potentials across trial events (the epoch period and set of 5 trial events are described in 4.6), resulting in a dimensionality of 251 time points (at 2 ms resolution) by 5 events per trial in the neural data. For participants with 2 recording sessions, we further concatenated the trials across sessions. This was done separately for each contact within an ROI, resulting in a per-subject design matrix of dimensionality: *nEvents* x *nContacts* x *nTimepoints* (251). To prepare the feature matrix and target vector for our decoding analyses, we split each participant’s data into cross-validation (CV) train/test folds, and separately concatenated the design matrices across all subjects for each fold. This ensured balance in the per-subject contribution to decoding, across CV folds. To create the final pseudo-population matrix, we concatenated across subjects (for each fold) along the *nEvents* and *nContact* dimensions for each ROI; this approach avoids the assumption that each trial occurs at the same time across participants and results in a largely sparse feature matrix (see Figure S7).

#### 4.10.3 Decoding model and procedure

The variable of interest to be decoded was binarized into to +1 and -1 around 0 (i.e. the mean) as the target vector for classification analyses. The target vector and feature matrix were split into a 10 folds for cross-validation (CV) along the *nEvents* dimension, and each fold was normalized (i.e. centered and scaled to unit variance) by computing statistics on the training set and applying to both training and test sets [76]. All feature preprocessing (see 4.10.1) and normalization was conducted prior to the between-subject concatenation described in 4.10.2. We specified logistic regression decoding models with L2 regularization to account for potentially high dimensionality in the feature matrix. The 10-fold CV decoding analysis was run at each time point in the *nTimepoints* dimension independently and the resulting decoding curve was submitted for statistical analyses (see 4.10.4. We employed the receiver operating characteristic (ROC) area under the curve (AUC) as our classification accuracy metric for increased robustness against cutoff values of the probabilistic predictions of our logistic classifier. Decoding curves were low-pass filtered (1st-order Butterworth filter at 0.1 Hz) for visualization; all statistics are conducted on non low-pass filtered data.

#### 4.10.4 Statistical analysis

We used a non-parametric maximum cluster statistic approach [77] to determine temporal clusters of significant decoding while keeping FWE < 0.05. Within a ROI and for a given computational model, we permuted the target vector labels 1000 times and tested the model on the permuted labels to simulate a null distribution of mean CV accuracy, at each timepoint. Each permutation’s shuffled mean CV accuracy was thresholded at the 95^th^ percentile of this distribution and the maximum above-threshold AUC of the resulting clusters was extracted; these comprised the max. cluster-statistic null distribution. The procedure was applied to the un-shuffled decoding curve and each above-threshold cluster was deemed significant if its cluster statistic was above the 95^th^ percentile of the max. cluster-statistic null distribution, keeping FWE < 0.05. Cluster-wide p-values reported in the main text are calculated as 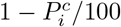 with 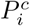 denoting the *i*^*th*^ percentile of cluster *c*’s above-threshold statistic along the max. cluster-statistic null distribution. Results in exploratory ROIs were FDR-corrected against the full set of 11 ROIs and corrected p-values are reported as *q*-values. To evaluate whether significant clusters differed in their latencies, we conducted non-parametric Mann-Whitney U-rank tests to compare the timings respective to each significant cluster, for pairs of temporally adjacent clusters as ordered by the latency of the peak decoding accuracy. The Mann-Whitney U-statistic and p-value of the test are reported alongside the [min,max] of the span of significant clusters.

#### 4.10.5 Feature importance

We used a leave-one-feature-out procedure for determining the contribution of each contact to the overall decoding accuracy for a given ROI and variable. The weight of a given feature (i.e. contact) was defined as the change in mean CV ROC AUC between a full model and one in which only that feature was dropped, and resulting features weights were of dimensionality *nContacts* x *nTimepoints* per ROI. To test for differences in the contribution of each OFC contact in decoding distinct variables, we z-scored the feature weights and averaged the normalized feature weights across the overlapping period of significant decoding across variables (OUT, RePE, and RiPE) per contact. We then computed Pearson’s correlation (r) values between the average normalized feature weights respective to each variable in a tested pair (i.e. OUT and RePE; RePE and RiPE) across contacts.

### 4.11 Decoding generalization

For our across-variable and temporal generalized decoding [18] analysis focusing on Anterior Insula and OFC, we applied the same feature preprocessing steps and leveraged the same pseudo-population design reported above. Following the procedure reported in section 4.10.3, we ran a 10-fold CV test using L2-regularized logistic regression classifiers and report ROC AUC. Here we trained the classifier on the binarized RePE target vector at each time-point, and without further tuning model weights tested the model on its ability to generalize on the out-of-sample binarized RiPE target vector across each time-point, with performance measured by the average across-fold ROC AUC. We constrained the time window for generalized decoding analysis to 0.200 - 0.500 sec after the onset of card 2, to encompass the time points in which we were able to decode RePE and RiPE alone (i.e., without generalization) in both ROIs and the a priori hypothesis that RePE (and thus the extent to which it can inform RiPE) can only occur after the onset of card 2 when the outcome is known. To test for robustness, we repeated this procedure without constraining the time window, thus using the entire epoch after the onset of card 2 (see Figure S3). For the OFC ROI, we also ran this procedure trained on Outcome and tested on RePE given our finding that OFC contacts significantly decode both these variables in overlapping periods after the onset of card 2. We applied the same max. cluster-statistic approach described in section 4.10.4 but extending the cluster based inference to two dimensions, and report results exceeding the 95^th^ and 99^th^ percentiles of the permuted null distribution, both keeping FWE < 0.05 at the cluster level. We report cluster-wide p-values with respect to the max. cluster-statistic null distribution.

## 5 Acknowledgments

This work was supported by a grant to J.P.O. and M.A.H. from NIMH (R01MH111425). We thank members of the Human Reward and Decision Making lab and the Human Brain Research Lab for discussions and feedback. We thank all participants and their families for participating in this research.

## 5.1 Contributions

V.M., J.C., O.F., P.G., M.S., C.K.K., H.K., and H.O. collected the data. J.C. and J.P.O. conceived of the experiment. V.M. and J.P.O. designed the data analysis. V.M. conducted the data analysis. V.M., J.C., and J.P.O. wrote the initial draft of the manuscript. All authors revised the manuscript.

