## Supplementary Material for "Temporally organized representations of reward and risk in the human brain"

### 1 Supplementary Materials

#### 1.1 Cross-correlation analysis

We investigated whether neural activity was correlated across regions of interest by running a shuffle-corrected cross-correlation analysis [?] on the neural activity between two regions, following the procedure described in [?] but as applied to iEEG data. We specifically examined whether the average cross-correlation between ROIs was moderated by our computational variables of interest. We followed the same feature preprocessing procedure described in section **??** to regress out of the neural data variables other than the current moderating variable of interest, and normalized the neural data for each contact. We then computed the cross-correlation between the activity in pairs of contacts between regions A and B recorded from the same participant according to:

$$C^r\tau = \sum_{-\infty}^{\infty} S_A^r(t) \cdot S_B^r(t+\tau) \equiv S_A^r \otimes S_B^r$$

where $S_A^r$, $S_B^r$ denotes the activity from contacts in ROIs A and B, respectively, across the relevant epoch on trial $r$. We defined the shuffle-corrected cross-correlogram as:

$$V = < S_A^r \otimes S_B^r > - < S_A^r > \otimes < S_B^r >$$

where the < > operator denotes averaging over trials, separately for across-trials splits according to low and high values of the computational variable of interest (following the same approach for binarization described in section **??**. This procedure of regularizing the cross-correlogram with the shuffle-correcting term is equivalent to permutation-based approaches (e.g. shift predictor; see [?, ?]) in correcting for time-locked covariation (e.g. caused by stimulus presentation) and reflected in each contact separately. In other words, the shuffle-corrector ensures that if $SA$ and $SB$ are independent, the expected value of V is zero. We transformed the resulting cross-correlations using Fisher's r-to-z transformation prior to averaging across pairs [?]. We summed the average z-values separately for high and low trials of the variable, and for negative and positive lag regions of the resulting cross-correlogram and tested for significant differences in the average cross-correlation between high and low levels of the computational variable of interest.

For each computational variable of interest, we defined pairs of ROIs to submit to the cross-correlation analysis based on regions in which we could decode each respective variable in our main results. For a given variable, we aligned the ROIs based on their peak decoding accuracy and tested for differences in the cross-correlation between adjacent regions (e.g. for outcome representations: Angular Gyrus and Hippocampus, Hippocampus and Amygdala, etc) as a function of trials sorted along high or low levels of the variable. Cross-correlation results for outcome representations are reported in the main text and in Figure S5; for all variables other than outcome, we did not find evidence of differences in the between-ROI cross-correlations as a function of the high versus low level of that computational variable.

#### 1.2 Contact coordinates

We provide the contact coordinates for our main *a priori* regions of interest including subregions of the prefrontal cortex, the amygdala, and the anterior insula. We also include coordinates for contacts in the posterior insula for comparison. Patient indices indicate contacts that come from the same individual. Coordinates are in CIT168toMNI transformed space. Adjacent contacts were paired for bipolar (BP) referencing and assigned with the label of anode (A) or cathode (C). Laterality denotes left (L) or right (R) contact location and implantation side. The Brodmann areas (BA) are provided for cortical coordinates when available.

#### 1.3 Supplementary Figures

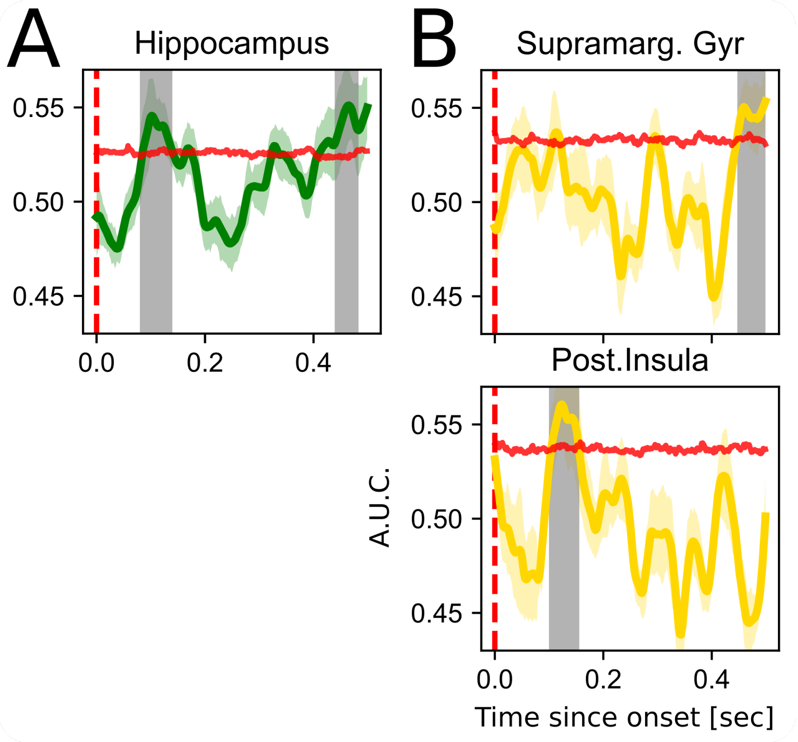

Supplementary Figure 1: Decoding of (A) expected value and (B) expected risk after the offset of card 1 in exploratory ROIs. Lines depict cross-validated receiver operating characteristic (ROC) accuracy (AUC), and shaded coloured areas show standard error in ROC accuracy across folds. Horizontal red lines depict the 95th percentile of the permuted null distribution at each time point, and periods of statistical significance are shown in the shaded grey region (cluster corrected FWE < 0.05). Decoding curves are low-pass filtered (1st-order Butterworth filter at 0.1 Hz) for visualization; all statistics are conducted on non low-pass filtered data.

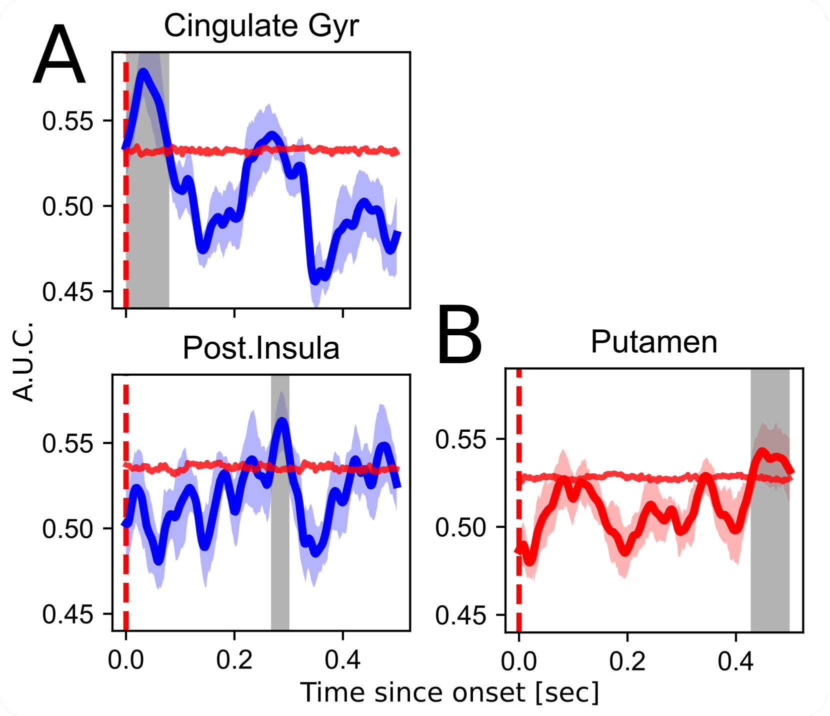

Supplementary Figure 2: Decoding of (A) reward prediction error and (B) risk prediction error in exploratory ROIs. Description follows that described in Figure S1.

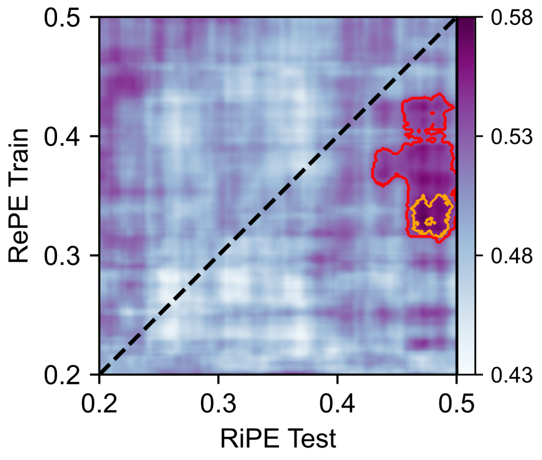

Supplementary Figure 3: Generalized decoding in anterior insula. Same as Figure **??**A, but across the entire epoch after the onset of card 2 [0.0 - 0.500 sec]. Significant decoding areas are highlighted by the red (p = 0.031) and orange (p = 0.003) contours (both cluster corrected FWE < 0.05).

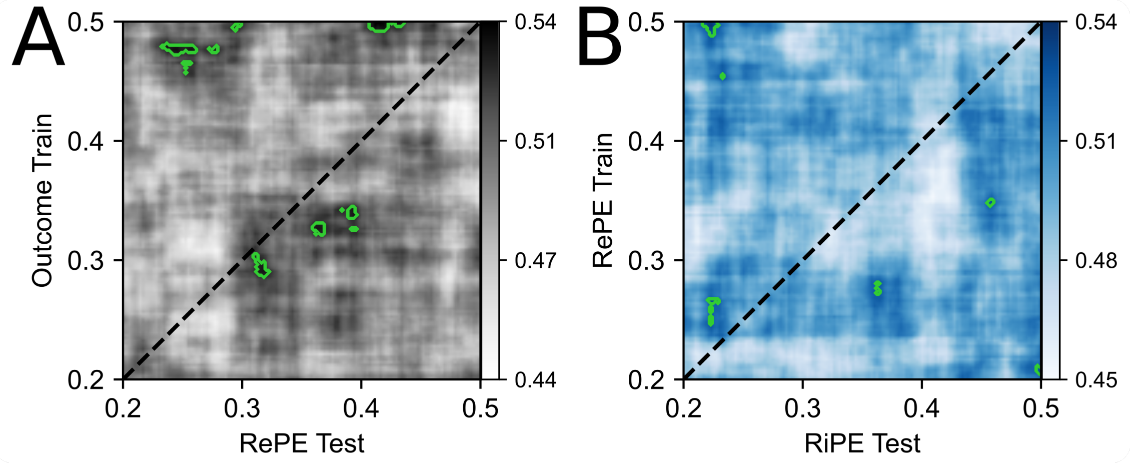

Supplementary Figure 4: Lack of generalized decoding in OFC for models (A) trained on the multivariate signal in OFC encoding outcome and tested on RePE, and (B) trained on RePE and tested on RiPE. Areas highlighted in green reflect decoding accuracies above the 95th percentile of the permuted null distribution, but for both models no cluster survived correction at FWE < 0.05.

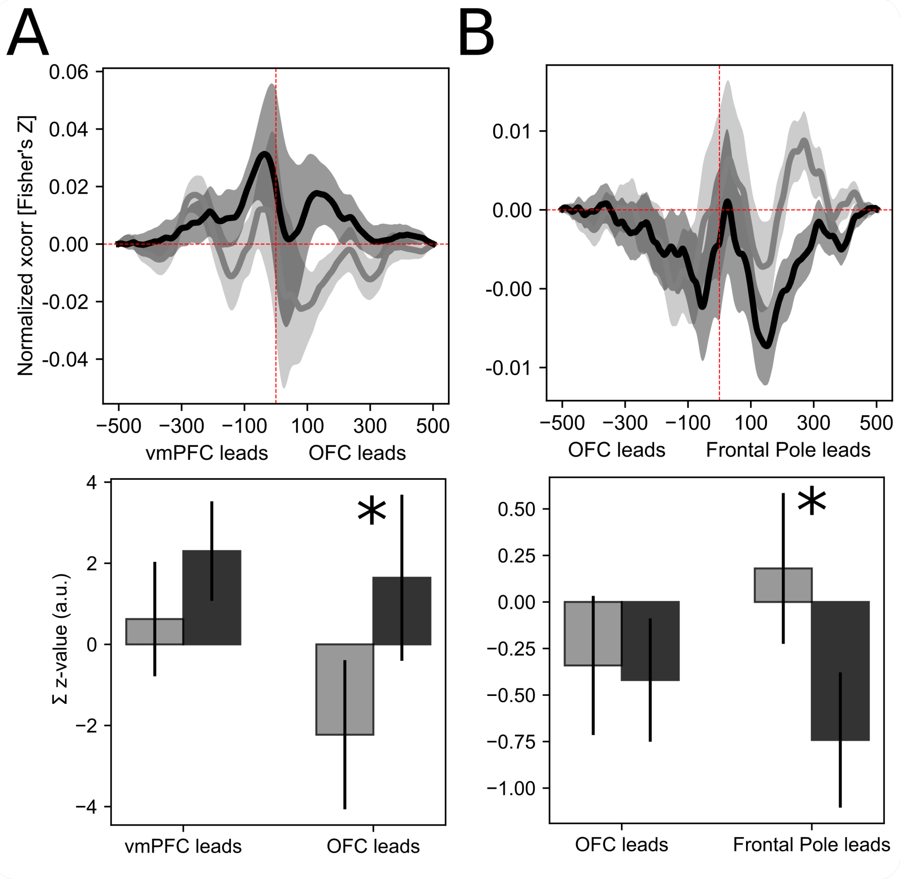

Supplementary Figure 5: Cross-correlation analysis between pairs of outcome-related prefrontal ROIs. A) Contacts in lateral OFC lead those in vmPFC as a function of outcome during the period ranging 0.500 sec after the onset of card 2. B) Contacts in frontal pole, which incorporate anterior OFC, lead those in lateral OFC. Top panels depict the shuffle-corrected cross-correlogram with traces and shaded areas depicting the mean and sem normalized cross-correlation over across-ROI contact pairs. The bottom panels depict a summary of the cross-correlogram calculated by integrating over the area under each mean trace, separately for low and high outcome trials and for negative and positive lags (* p < 0.05).

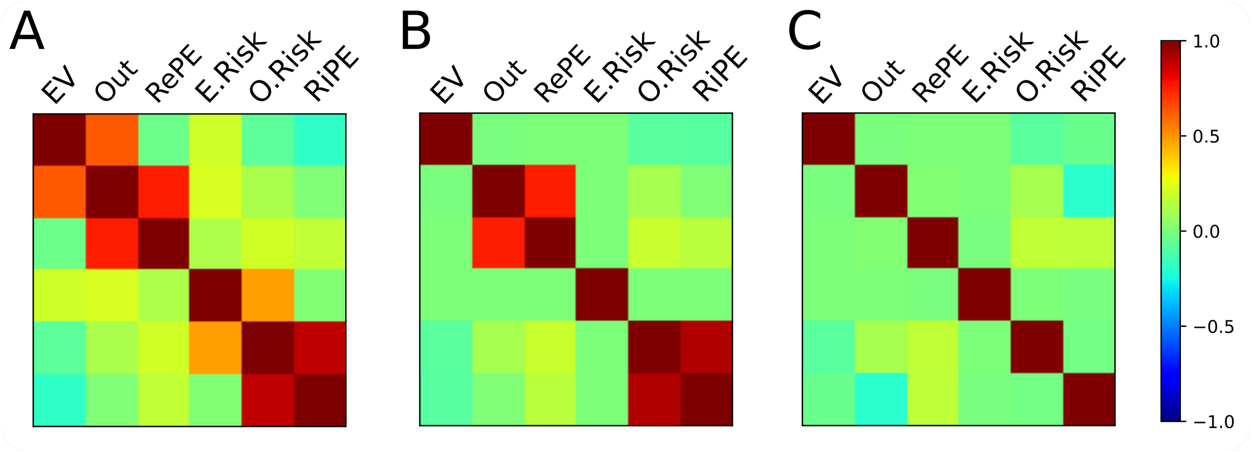

Supplementary Figure 6: Correlation between computational variables. (A) As derived directly from behavior and the normative model, there are substantial correlations between the computational variables. (B) Our expanded design matrix spaced out comptuational variables across time within a trial and situated each variable at their hypothesized occurence given the information available to the participant at that moment (e.g. outcome after the second card is presented). This approach decoupled most variables. (C) Variables that remained highly correlated after our expanded design matrix (i.e., outcome and RePE; observed risk and RiPE) were statistically orthogonalized such that the outcome and observed risk variables retained shared variance with RePE and RiPE, respectively.

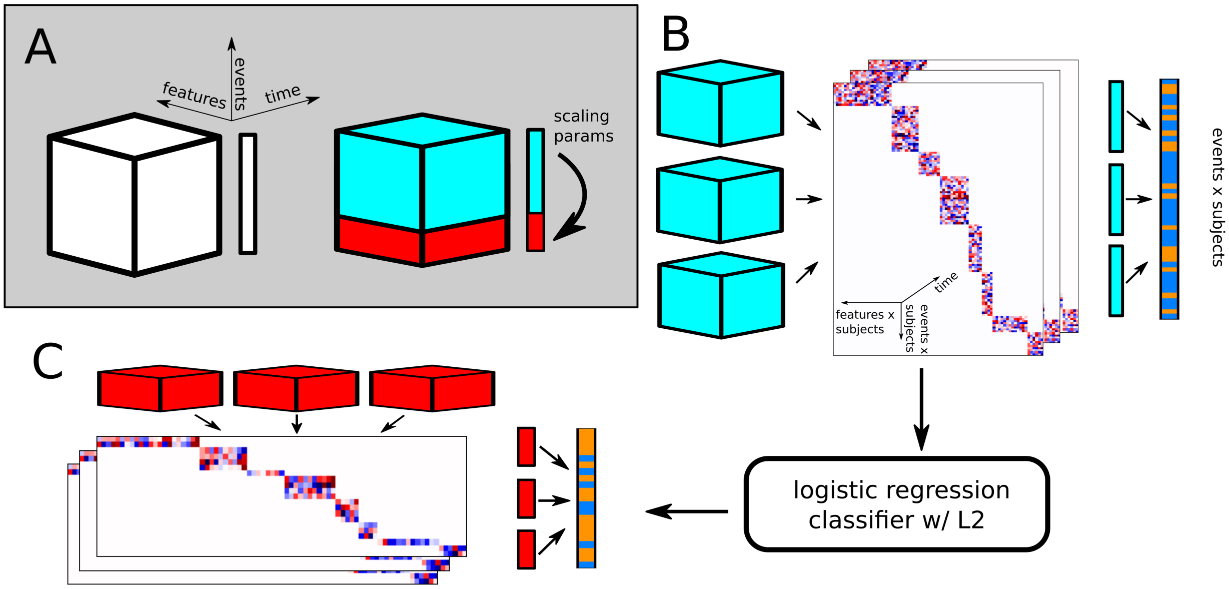

Supplementary Figure 7: Overview of pseudo-population decoding approach. A) Left: The features matrix for a single participant is organized along features (neural sources), events (within-trial events) and time (samples) dimensions. The target vector has the dimensionality of events. Right: The features are preprocessed and split into train (cyan) and test (red) folds within-subject. Preprocessing parameters are estimated on the training data and applied to both the train and test sets. B) The train sets across participants are concatenated diagonally into a pseudo-population matrix, which has features x subjects, events x subjects and time dimensions, for a given ROI. Each colored tile in the pseudo-population concatenated matrix (middle) corresponds to the neural data from one participant. The target vectors across participants are concatenated and of events x subjects length. The concatenated feature matrix and target vector are used to train the logistic regression classifier, which is then tested on the left-out data (C) which is organized across participants similarly. A single cross-validation fold is depicted; this procedure is iterated across 10 folds and mean receiver operating characteristic area-under-the curve (ROC AUC) metrics are reported as the accuracy metric.

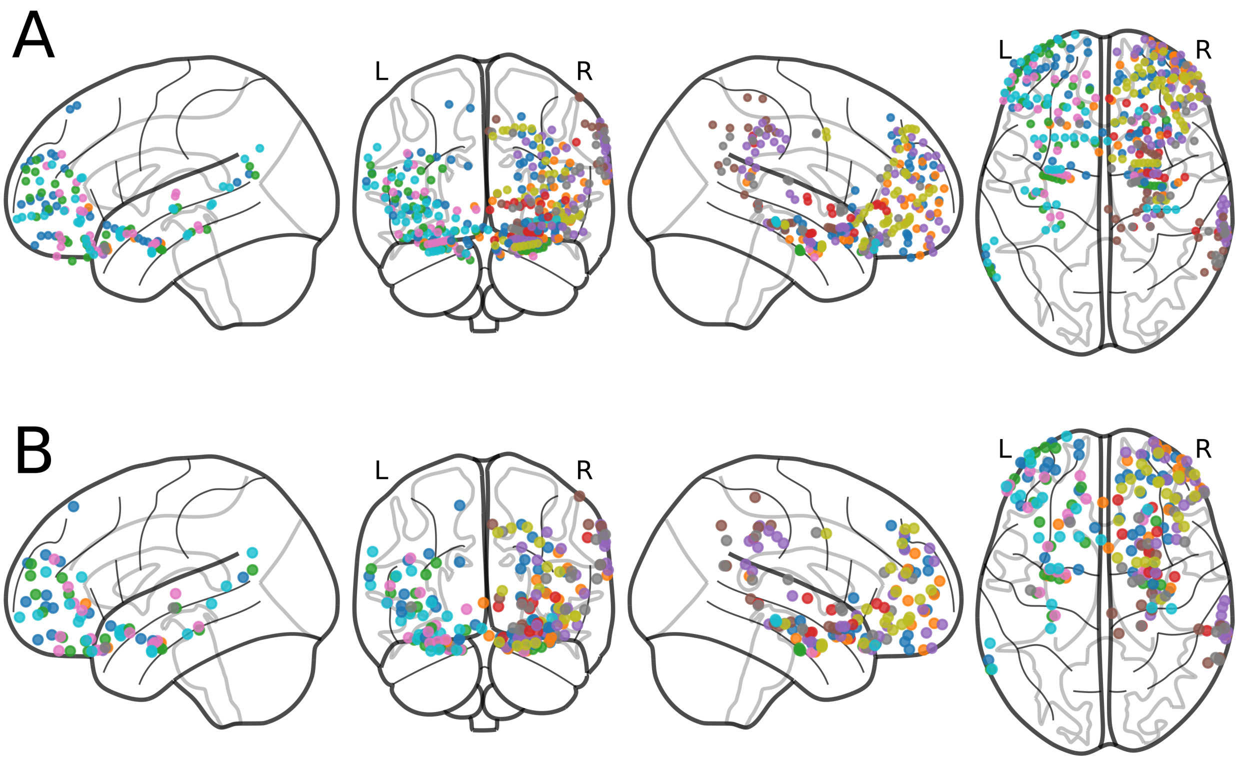

Supplementary Figure 8: Contact coordinates in original recording (A) and projected bipolar source (B) locations displayed over template space. Bipolar locations were interpolated by averaging across anode and cathode pair coordinates across three dimensions. Panel B is the same as Figure **??**B, but organized by participant rather than ROIs. Different colors identify distinct participants.

| Patient Idx | X | Y | Z | Source Idx | BP label | Laterality | BA |
|---|---|---|---|---|---|---|---|
| 0 | 9.17 | 50.07 | -30.11 | 0 | A | R | – |
| 0 | 18.26 | 48.41 | -21.85 | 0 | C | R | 11 |
| 0 | 27.06 | 48.56 | -18.47 | 1 | A | R | 10 |
| 0 | 39.91 | 48.42 | -17.46 | 1 | C | R | 10 |
| 0 | -7.49 | 42.45 | 53.05 | 2 | A | L | 8 |
| 0 | -19.42 | 38.6 | 55.18 | 2 | C | L | – |
| 0 | -8.8 | 67.92 | 20.62 | 3 | A | L | 10 |
| 0 | -18.28 | 64.44 | 24.19 | 3 | C | L | 10 |
| 0 | -26.37 | 59.27 | 25.72 | 4 | A | L | 10 |
| 0 | -35.02 | 52.95 | 28.3 | 4 | C | L | 10 |
| 0 | -25.6 | 69.79 | -0.23 | 5 | A | L | – |
| 0 | -31.81 | 64.61 | 1.76 | 5 | C | L | 10 |
| 0 | -40.97 | 59.39 | 4.04 | 6 | A | L | – |
| 0 | -48.12 | 51.92 | 4.77 | 6 | C | L | 46 |
| 0 | 8.12 | 45.84 | -26.73 | 7 | A | R | 11 |
| 0 | 11.27 | 47.66 | -16.54 | 7 | C | R | 11 |
| 0 | 19.1 | 49.59 | 14.72 | 8 | A | R | 10 |
| 0 | 22.35 | 48.89 | 22.67 | 8 | C | R | 10 |
| 0 | 21.35 | 39.41 | 38.45 | 9 | A | R | 8 |
| 0 | 21.11 | 38.76 | 48.39 | 9 | C | R | 8 |
| 0 | -21.52 | 49.94 | -1.17 | 10 | A | L | – |
| 0 | -32.05 | 54 | -4.3 | 10 | C | L | 10 |
| 0 | -42.23 | 56.03 | -3.92 | 11 | A | L | 10 |
| 0 | -49.94 | 56.15 | -0.35 | 11 | C | L | – |
| 0 | -42.86 | 45.5 | 19.85 | 12 | A | L | 10 |
| 0 | -41.82 | 50.58 | 26.73 | 12 | C | L | – |
| 0 | 28.4 | 62.84 | 15.13 | 13 | A | R | 10 |
| 0 | 32.8 | 56.82 | 21.23 | 13 | C | R | – |
| 0 | 35.37 | 50.39 | 26.95 | 14 | A | R | 10 |
| 0 | 36.78 | 42.63 | 34.18 | 14 | C | R | 9 |
| 0 | 20.31 | 62.8 | 21.73 | 15 | A | R | 10 |
| 0 | 25.81 | 56.78 | 25.56 | 15 | C | R | 10 |
| 0 | 27.63 | 50.07 | 32.41 | 16 | A | R | 9 |
| 0 | 27.06 | 43.21 | 40.27 | 16 | C | R | 9 |
| 0 | 37.5 | 61.11 | -7.17 | 17 | A | R | – |
| 0 | 43.66 | 57.02 | -0.35 | 17 | C | R | 10 |
| 0 | 31.15 | 65.93 | 1.96 | 18 | A | R | 10 |
| 0 | 37.15 | 60.35 | 9.04 | 18 | C | R | – |
| 0 | 43.35 | 55.67 | 13.76 | 19 | A | R | – |
| 0 | 47.42 | 49.73 | 20.15 | 19 | C | R | – |
| 0 | -8.5 | 62.21 | -23.17 | 20 | A | L | – |
| 0 | -18.56 | 58.71 | -17.89 | 20 | C | L | 11 |
| 0 | -27.07 | 54.72 | -17.56 | 21 | A | L | 10 |
| 0 | -38.32 | 50.59 | -18.02 | 21 | C | L | – |
| 1 | 37.91 | 46.49 | -0.75 | 22 | A | R | 10 |
| 1 | 46.08 | 50.51 | 3.51 | 22 | C | R | 10 |
| 1 | 9.43 | 55.23 | -28.78 | 23 | A | R | – |
| 1 | 17.62 | 59.26 | -23.66 | 23 | C | R | – |
| 1 | 29.29 | 61.4 | -17.52 | 24 | A | R | – |
| 1 | 33.86 | 64.94 | -5.08 | 24 | C | R | – |
| 1 | 40.14 | 53.34 | -13.82 | 25 | A | R | 10 |
| 1 | 48.53 | 47.21 | -10.47 | 25 | C | R | 10 |
| 1 | 40.05 | 60.46 | -6.8 | 26 | A | R | 10 |
| 1 | 47.69 | 54.91 | -2.67 | 26 | C | R | – |
| 1 | 52.65 | 46.66 | 2.45 | 27 | A | R | 46 |
| 1 | 56.27 | 36.96 | 11.52 | 27 | C | R | 46 |
| 1 | 31.04 | 65.83 | 0.69 | 28 | A | R | 10 |

**Table 1 continued from previous page**

| Patient Idx | X | Y | Z | Source Idx | BP label | Laterality | BA |
|---|---|---|---|---|---|---|---|
| 1 | 39.33 | 60.35 | 8.88 | 28 | C | R | – |
| 1 | 45.64 | 52.83 | 15.42 | 29 | A | R | – |
| 1 | 48.34 | 43.95 | 22.39 | 29 | C | R | 10 |
| 1 | 26.35 | 68.73 | 8.63 | 30 | A | R | – |
| 1 | 33.05 | 63.73 | 16.08 | 30 | C | R | – |
| 1 | 39.4 | 56.51 | 23.73 | 31 | A | R | – |
| 1 | 44.37 | 48.2 | 31.91 | 31 | C | R | – |
| 2 | -7.58 | 41.25 | -32.01 | 32 | A | L | – |
| 2 | -17.01 | 43.49 | -21.65 | 32 | C | L | 11 |
| 2 | -26.84 | 44.97 | -20.08 | 33 | A | L | 11 |
| 2 | -37.15 | 48.52 | -18.34 | 33 | C | L | – |
| 2 | -23.08 | 67.03 | 19.23 | 34 | A | L | – |
| 2 | -30.21 | 61.4 | 24.65 | 34 | C | L | – |
| 2 | -29.51 | 67.26 | 12.77 | 35 | A | L | – |
| 2 | -35.18 | 61.95 | 17.96 | 35 | C | L | – |
| 2 | -33.86 | 65.04 | 3.37 | 36 | A | L | – |
| 2 | -42.43 | 58.41 | 6.32 | 36 | C | L | – |
| 2 | -47.64 | 50.82 | 11.22 | 37 | A | L | – |
| 2 | -51.53 | 43 | 17.03 | 37 | C | L | – |
| 2 | -38.06 | 61.28 | -7.24 | 38 | A | L | – |
| 2 | -43.52 | 54.44 | -2.88 | 38 | C | L | 10 |
| 2 | -49.53 | 46.85 | 2.24 | 39 | A | L | 46 |
| 2 | -54.67 | 39.37 | 7.94 | 39 | C | L | 46 |
| 4 | 49.15 | 37.18 | -9.5 | 40 | A | R | 47 |
| 4 | 55.75 | 36.71 | -16.31 | 40 | C | R | – |
| 4 | 27.49 | 66.79 | 20.2 | 41 | A | R | – |
| 4 | 32.5 | 59.66 | 26.65 | 41 | C | R | – |
| 4 | 40.32 | 51.54 | 29.06 | 42 | A | R | – |
| 4 | 46.4 | 41.64 | 30.77 | 42 | C | R | – |
| 4 | 18.95 | 64.22 | 27.01 | 43 | A | R | – |
| 4 | 23.68 | 56.98 | 34.38 | 43 | C | R | 9 |
| 4 | 32.66 | 50.44 | 37.39 | 44 | A | R | – |
| 4 | 38.15 | 42.43 | 40.87 | 44 | C | R | – |
| 4 | 7.69 | 62.55 | -27.11 | 45 | A | R | – |
| 4 | 17.6 | 61.11 | -23.27 | 45 | C | R | – |
| 4 | 28.76 | 60.29 | -19.82 | 46 | A | R | – |
| 4 | 40.32 | 57.31 | -13.52 | 46 | C | R | 10 |
| 4 | 23.28 | 69.66 | -14.29 | 47 | A | R | – |
| 4 | 33.16 | 65.73 | -9.67 | 47 | C | R | – |
| 4 | 43.46 | 60.82 | -5.67 | 48 | A | R | – |
| 4 | 50.14 | 52.8 | -1.17 | 48 | C | R | – |
| 4 | 54.8 | 44.56 | 2.67 | 49 | A | R | 46 |
| 4 | 58.04 | 35.91 | 6.96 | 49 | C | R | 45 |
| 4 | 49.05 | 50.89 | 13.55 | 50 | A | R | – |
| 4 | 54.07 | 41.12 | 16.85 | 50 | C | R | – |
| 6 | -9.38 | 49.9 | -29.3 | 51 | A | L | – |
| 6 | -16.36 | 47.54 | -23.97 | 51 | C | L | 11 |
| 6 | -27.04 | 47.22 | -20.06 | 52 | A | L | – |
| 6 | -38.72 | 48.62 | -17.5 | 52 | C | L | 10 |
| 6 | -39.22 | 55.54 | 21.32 | 53 | A | L | – |
| 6 | -42.83 | 47.14 | 27.79 | 53 | C | L | 10 |
| 6 | -50.74 | 48.41 | 6 | 54 | A | L | – |
| 6 | -54.22 | 38.79 | 12.19 | 54 | C | L | – |
| 6 | -51.84 | 45.26 | -3.54 | 55 | A | L | 47 |
| 6 | -55.8 | 35.35 | 3.31 | 55 | C | L | 45 |
| 7 | 44.81 | 33.09 | 14.41 | 56 | A | R | 46 |

**Table 1 continued from previous page**

| Patient Idx | X | Y | Z | Source Idx | BP label | Laterality | BA |
|---|---|---|---|---|---|---|---|
| 7 | 53.88 | 37.44 | 19.92 | 56 | C | R | – |
| 7 | 35.96 | 36.88 | -21.98 | 57 | A | R | 47 |
| 7 | 46.92 | 35.93 | -18.35 | 57 | C | R | 47 |
| 7 | 55.01 | 43.12 | -8.1 | 58 | A | R | – |
| 7 | 58.26 | 34.92 | -0.58 | 58 | C | R | 45 |
| 7 | 46.35 | 55.22 | 9.87 | 59 | A | R | – |
| 7 | 51.14 | 46.1 | 17.74 | 59 | C | R | – |
| 8 | 36.45 | 36.79 | 15.81 | 60 | A | R | – |
| 8 | 34.67 | 40.61 | 23.52 | 60 | C | R | 9 |
| 8 | 32.11 | 43.86 | 30.76 | 61 | A | R | 9 |
| 8 | 30.5 | 46.62 | 36.13 | 61 | C | R | 9 |
| 8 | 13.3 | 45.41 | 40.63 | 62 | A | R | 9 |
| 8 | 17.44 | 48.37 | 42.19 | 62 | C | R | 9 |
| 8 | 21.93 | 50.74 | 42.16 | 63 | A | R | 9 |
| 8 | 26.84 | 52.8 | 41.6 | 63 | C | R | – |
| 8 | 37.28 | 34.86 | -11.08 | 64 | A | R | 47 |
| 8 | 43.08 | 36.48 | -10.6 | 64 | C | R | 47 |
| 8 | 49.06 | 37.23 | -9.46 | 65 | A | R | 47 |
| 8 | 53.77 | 37.46 | -6.36 | 65 | C | R | 47 |
| 8 | 17.29 | 39.37 | -16.12 | 66 | A | R | 11 |
| 8 | 23.7 | 40.68 | -14.01 | 66 | C | R | 11 |
| 8 | 28.41 | 43.54 | -12.29 | 67 | A | R | 47 |
| 8 | 32.98 | 46.57 | -11.57 | 67 | C | R | 47 |
| 8 | 39.44 | 48.12 | -10.27 | 68 | A | R | 10 |
| 8 | 45.23 | 49.01 | -9.19 | 68 | C | R | 10 |
| 8 | 49.24 | 49.3 | -8.61 | 69 | A | R | 10 |
| 8 | 53.12 | 49.4 | -7.24 | 69 | C | R | – |
| 8 | 19.72 | 53.62 | 0.64 | 70 | A | R | – |
| 8 | 23.29 | 56.84 | 3.53 | 70 | C | R | 10 |
| 8 | 28.31 | 60.21 | 6.33 | 71 | A | R | 10 |
| 8 | 33.78 | 62.74 | 8.31 | 71 | C | R | 10 |
| 8 | 39.43 | 47.85 | 16.28 | 72 | A | R | 10 |
| 8 | 45.59 | 46.83 | 17.57 | 72 | C | R | 10 |
| 9 | -46.34 | 35.55 | 14.78 | 73 | A | L | 46 |
| 9 | -51.54 | 36.41 | 14.42 | 73 | C | L | 46 |
| 9 | -15.69 | 72.19 | -8.47 | 74 | A | L | – |
| 9 | -25.21 | 69.34 | -7.39 | 74 | C | L | – |
| 9 | -36.72 | 64.07 | -6.69 | 75 | A | L | – |
| 9 | -46.14 | 56.9 | -6.57 | 75 | C | L | – |
| 9 | -43.64 | 37.15 | -10.72 | 76 | A | L | 47 |
| 9 | -49.07 | 38.31 | -10.24 | 76 | C | L | 47 |
| 9 | -38.62 | 59.38 | 12.84 | 77 | A | L | 10 |
| 9 | -42.31 | 52.58 | 22.07 | 77 | C | L | – |
| 9 | -53.22 | 45.83 | -3.88 | 78 | A | L | 47 |
| 9 | -55.16 | 39.58 | 2.87 | 78 | C | L | 45 |

Supplementary Table 1: Frontal Pole

| Patient Idx | X | Y | Z | Source Idx | BP label | Laterality | BA |
|---|---|---|---|---|---|---|---|
| 0 | 6.93 | 22.64 | -26.14 | 0 | A | R | 11 |
| 0 | 11.82 | 27.05 | -13.5 | 0 | C | R | 11 |
| 0 | -0.25 | 17.44 | -13.72 | 1 | A | L | – |
| 0 | -9.12 | 21.83 | -11.82 | 1 | C | L | – |
| 1 | -2.35 | 6.43 | -19.17 | 2 | A | R | – |
| 1 | 8.29 | 10.19 | -18.6 | 2 | C | R | – |
| 2 | -4.36 | 11.54 | -15.77 | 3 | A | L | 25 |
| 2 | -15.84 | 11.07 | -15.69 | 3 | C | L | – |
| 3 | 5.97 | 15.18 | -20.88 | 4 | A | R | 25 |
| 3 | 15.46 | 16.92 | -18.73 | 4 | C | R | 11 |
| 9 | 2.45 | 12.61 | -15.14 | 5 | A | L | |
| 9 | -4.78 | 13.04 | -14.48 | 5 | C | L | 25 |

Supplementary Table 2: Subcallosal Cortex

| Patient Idx | X | Y | Z | Source Idx | BP label | Laterality | BA |
|---|---|---|---|---|---|---|---|
| 0 | 32.84 | 28.73 | -24.08 | 0 | A | R | 47 |
| 0 | 41.47 | 32.4 | -21.28 | 0 | C | R | 47 |
| 0 | 14.54 | 14.99 | -22.65 | 1 | A | R | 11 |
| 0 | 18.5 | 16.42 | -12.63 | 1 | C | R | – |
| 0 | 31.38 | 18.32 | -19.16 | 2 | A | R | 47 |
| 0 | 36.1 | 20.78 | -7.75 | 2 | C | R | 13 |
| 0 | 10.63 | 5.77 | -17.7 | 3 | A | R | – |
| 0 | 21.13 | 7.66 | -17.42 | 3 | C | R | – |
| 0 | -41.25 | 32.55 | -4.05 | 4 | A | L | 47 |
| 0 | -50.74 | 31.28 | -7.05 | 4 | C | L | 47 |
| 1 | 18.88 | 12.7 | -20.22 | 5 | A | R | – |
| 1 | 32.02 | 15.74 | -20.19 | 5 | C | R | 47 |
| 1 | -29.79 | 21.78 | -24 | 6 | A | R | 47 |
| 1 | -20.34 | 21.22 | -25.9 | 6 | C | R | 47 |
| 1 | 32.89 | 28.94 | -24.51 | 7 | A | R | 47 |
| 1 | 40.49 | 30.32 | -20.19 | 7 | C | R | 47 |
| 2 | -12.54 | 32.69 | -29.89 | 8 | A | L | – |
| 2 | -22.38 | 31.91 | -24.13 | 8 | C | L | 11 |
| 2 | -32.51 | 33.1 | -22.87 | 9 | A | L | 47 |
| 2 | -45.31 | 34.47 | -19.6 | 9 | C | L | 47 |
| 2 | 35.33 | 24.41 | -21.62 | 10 | A | L | 47 |
| 2 | 23.94 | 22.08 | -26.78 | 10 | C | L | 47 |
| 2 | -30.71 | 23.35 | -23.01 | 11 | A | L | 47 |
| 2 | -40.6 | 22.23 | -17.25 | 11 | C | L | 47 |
| 3 | 22.89 | 31.5 | 0.03 | 12 | A | R | – |
| 3 | 32.12 | 30.35 | -0.68 | 12 | C | R | 45 |
| 4 | -25.12 | 24.34 | -24.69 | 13 | A | R | 47 |
| 4 | -13.31 | 23.87 | -28.24 | 13 | C | R | – |
| 4 | 19.36 | 26.57 | -28.51 | 14 | A | R | – |
| 4 | 28.51 | 29.62 | -23.33 | 14 | C | R | 47 |
| 4 | 40 | 32.5 | -19.41 | 15 | A | R | 47 |
| 4 | 50.47 | 31.98 | -17.77 | 15 | C | R | 47 |
| 4 | 15.86 | 7.86 | -20.79 | 16 | A | R | – |
| 4 | 25.08 | 13.7 | -16.16 | 16 | C | R | – |
| 4 | 36.33 | 18.65 | -8.87 | 17 | A | R | 13 |

**Table 3 continued from previous page**

| Patient Idx | X | Y | Z | Source Idx | BP label | Laterality | BA |
|---|---|---|---|---|---|---|---|
| 4 | 46.57 | 20.67 | -5.27 | 17 | C | R | – |
| 5 | 20.79 | 10.51 | -14.83 | 18 | A | R | – |
| 5 | 22.67 | 9.15 | -14.58 | 18 | C | R | – |
| 5 | 24.7 | 7.68 | -14.2 | 19 | A | R | – |
| 5 | 26.75 | 6.26 | -13.64 | 19 | C | R | – |
| 6 | 27.56 | 24.59 | -23.82 | 20 | A | L | 47 |
| 6 | 17.04 | 23.26 | -26 | 20 | C | L | 11 |
| 6 | -14.44 | 25.59 | -26.88 | 21 | A | L | 11 |
| 6 | -25.23 | 28.71 | -25.8 | 21 | C | L | – |
| 6 | -36.61 | 30.45 | -22.25 | 22 | A | L | 47 |
| 6 | -47.6 | 31.39 | -17.68 | 22 | C | L | 47 |
| 7 | -23.84 | 23.11 | -23.73 | 23 | A | R | 47 |
| 7 | -13.25 | 22.28 | -25.93 | 23 | C | R | 11 |
| 7 | 15.86 | 24.07 | -26.55 | 24 | A | R | 11 |
| 7 | 24.11 | 26.31 | -23.67 | 24 | C | R | 47 |
| 7 | 33.52 | 27.95 | -20.58 | 25 | A | R | 47 |
| 7 | 46.46 | 29.26 | -15.55 | 25 | C | R | 47 |
| 8 | 45.78 | 18.93 | -10.67 | 26 | A | R | 47 |
| 8 | 44.89 | 21.68 | -7.99 | 26 | C | R | 13 |
| 8 | 43.2 | 23.93 | -5 | 27 | A | R | 13 |
| 8 | 40.99 | 26.96 | -1.71 | 27 | C | R | 47 |
| 8 | 6.42 | 23.04 | -23.82 | 28 | A | R | 11 |
| 8 | 10.99 | 24.2 | -21.02 | 28 | C | R | 11 |
| 9 | 29.97 | 30.38 | -21.74 | 29 | A | L | 47 |
| 9 | 20.11 | 29.45 | -25.54 | 29 | C | L | 11 |
| 9 | -9.98 | 29.13 | -28.33 | 30 | A | L | – |
| 9 | -20.25 | 31.38 | -25.6 | 30 | C | L | 11 |
| 9 | -31.35 | 33.12 | -22.25 | 31 | A | L | 47 |
| 9 | -42.78 | 33.68 | -19.65 | 31 | C | L | 47 |
| 9 | -10.5 | 14.36 | -15.29 | 32 | A | L | 11 |
| 9 | -16.5 | 14.73 | -15.57 | 32 | C | L | 11 |
| 9 | -21.35 | 14.53 | -16.58 | 33 | A | L | – |
| 9 | -27.16 | 14.63 | -18.16 | 33 | C | L | 13 |
| 9 | -33.57 | 14.1 | -18.71 | 34 | A | L | 13 |
| 9 | -38.52 | 13.87 | -17.91 | 34 | C | L | – |
| 9 | -31 | 36.59 | -5.99 | 35 | A | L | 47 |
| 9 | -37.02 | 36.38 | -9.15 | 35 | C | L | 47 |

Supplementary Table 3: Frontal Orbital Cortex

| Patient Idx | X | Y | Z | Source Idx | BP label | Laterality | BA |
|---|---|---|---|---|---|---|---|
| 0 | 23.18 | -10.44 | -11.2 | 0 | A | R | – |
| 0 | 32.88 | -13.29 | -8.52 | 0 | C | R | – |
| 0 | 24.9 | -2.59 | -23.63 | 1 | A | R | – |
| 0 | 30.3 | -0.81 | -23.34 | 1 | C | R | – |
| 0 | -9.54 | -6.6 | -20.96 | 2 | A | L | – |
| 0 | -18.12 | -1.08 | -21.2 | 2 | C | L | – |
| 0 | -28.4 | 1.41 | -20.71 | 3 | A | L | – |
| 0 | -36.92 | 4.1 | -19.2 | 3 | C | L | – |
| 3 | 24.64 | -5.05 | -16.25 | 4 | A | R | – |
| 3 | 26.77 | -5.58 | -16.44 | 4 | C | R | – |
| 3 | 28.93 | -6.18 | -16.61 | 5 | A | R | – |
| 3 | 31.27 | -6.83 | -16.66 | 5 | C | R | – |
| 4 | 28.11 | -5.81 | -21.37 | 6 | A | R | – |
| 4 | 30.62 | -5.4 | -21.96 | 6 | C | R | – |
| 5 | 28.72 | 4.91 | -13.25 | 7 | A | R | – |
| 5 | 30.61 | 3.56 | -13.03 | 7 | C | R | – |
| 6 | -24.63 | -5.48 | -21.86 | 8 | A | L | – |
| 6 | -26.88 | -4.81 | -22.37 | 8 | C | L | – |
| 6 | -29.1 | -4.15 | -22.83 | 9 | A | L | – |
| 6 | -31.28 | -3.5 | -23.29 | 9 | C | L | – |
| 7 | 15.66 | -10.95 | -14.22 | 10 | A | R | – |
| 7 | 17.54 | -10.42 | -14.73 | 10 | C | R | – |
| 8 | 17.7 | 0.04 | -24.71 | 11 | A | R | – |
| 8 | 19.99 | 0.22 | -24.45 | 11 | C | R | – |
| 8 | 22.39 | 0.39 | -24.12 | 12 | A | R | – |
| 8 | 24.99 | 0.39 | -23.65 | 12 | C | R | – |
| 8 | 27.64 | 0.3 | -23.15 | 13 | A | R | – |
| 8 | 30.2 | 0.23 | -22.71 | 13 | C | R | – |
| 9 | -23.78 | -3.15 | -26.36 | 14 | A | L | – |
| 9 | -26.24 | -3.34 | -26.95 | 14 | C | L | – |
| 9 | -28.41 | -3.46 | -27.26 | 15 | A | L | – |
| 9 | -30.69 | -3.25 | -27.62 | 15 | C | L | – |

Supplementary Table 4: Amygdala

| Patient Idx | X | Y | Z | Source Idx | BP label | Laterality | BA |
|---|---|---|---|---|---|---|---|
| 0 | 30.43 | 11.03 | -15.55 | 0 | A | R | – |
| 0 | 42.36 | 15.44 | -12.89 | 0 | C | R | 13 |
| 2 | -25.02 | 10.06 | -14.99 | 1 | A | L | – |
| 2 | -33.81 | 8.41 | -13.92 | 1 | C | L | 13 |
| 3 | 24.31 | 18.42 | -16.08 | 2 | A | R | – |
| 3 | 36.03 | 16.42 | -7.84 | 2 | C | R | 13 |
| 4 | 26.94 | 10.13 | -13.37 | 3 | A | R | – |
| 4 | 32.44 | 11.4 | -12.81 | 3 | C | R | – |
| 4 | 38.05 | 12.51 | -12.78 | 4 | A | R | 13 |
| 4 | 44.17 | 13.54 | -13.39 | 4 | C | R | – |
| 6 | -25.71 | 7.27 | -16.46 | 5 | A | L | – |
| 6 | -36.34 | 10.79 | -16.8 | 5 | C | L | 13 |
| 7 | 40.47 | 10.66 | -4.76 | 6 | A | R | 13 |
| 7 | 51 | 12.66 | -3.66 | 6 | C | R | – |

Supplementary Table 5: Anterior Insula

| Patient Idx | X | Y | Z | Source Idx | BP label | Laterality | BA |
|---|---|---|---|---|---|---|---|
| 2 | -34.94 | -16.07 | -3.39 | 0 | A | L | – |
| 2 | -39.61 | -17.92 | -1.97 | 0 | C | L | 13 |
| 3 | 34.38 | -8.04 | 3.97 | 1 | A | R | – |
| 3 | 45.73 | -7.6 | 1.48 | 1 | C | R | 13 |
| 6 | -33.09 | -16.91 | 6.22 | 2 | A | L | – |
| 6 | -38.18 | -16.41 | 4.38 | 2 | C | L | 13 |
| 7 | 32.23 | -19.32 | 13.57 | 3 | A | R | 13 |
| 7 | 37.62 | -17.98 | 12.77 | 3 | C | R | 13 |
| 9 | -37.49 | -15.75 | -2.92 | 4 | A | L | – |
| 9 | -43.9 | -15.03 | -2.55 | 4 | C | L | 22 |

Supplementary Table 6: Posterior Insula
